# Constitutive Plasma Membrane Turnover in T-REx293 cells via Ordered Membrane Domain Endocytosis under Mitochondrial Control

**DOI:** 10.1101/2024.01.17.576124

**Authors:** Christine Deisl, Orson W. Moe, Donald W. Hilgemann

**Affiliations:** Department of Physiology, University of Texas Southwestern Medical Center, Dallas, Texas, USA; Department of Internal Medicine, University of Texas Southwestern Medical Center, Dallas, Texas, USA; Charles and Jane Pak Center for Mineral Metabolism and Clinical Research, University of Texas Southwestern Medical Center, Dallas, Texas, USA

## Abstract

Clathrin/dynamin-independent endocytosis of ordered plasma membrane domains (**<u>o</u>**rdered **<u>m</u>**embrane **<u>d</u>**omain **<u>e</u>**ndocytosis, OMDE) can become massive in response to cytoplasmic Ca elevations, G protein activation by non-hydrolyzable GTP analogs, and enhanced oxidative metabolism. In patch-clamped murine bone marrow macrophages (BMMs), cytoplasmic succinate and pyruvate, but not β-hydroxybutyrate, induce OMDE of 75% of the plasma membrane within 2 min. The responses require palmitoylation of membrane proteins, being decreased by 70% in BMMs lacking the acyltransferase, DHHC5, by treatment with carnitine to shift long-chain acyl groups from cytoplasmic to mitochondrial acyl-CoAs, by bromopalmitate/albumin complexes to block DHHCs, and by the mitochondria-specific cyclosporin, NIM811, to block permeability transition pores that may release mitochondrial coenzyme A into the cytoplasm. Using T-REx293 cells, OMDE amounts to 40% with succinate, pyruvate, or GTPγS, and it is inhibited by actin cytoskeleton disruption. Pyruvate-induced OMDE is blocked by the hydrophobic antioxidant, edaravone, which prevents permeability transition pore openings. Using fluorescent 3kD dextrans to monitor endocytosis, OMDE appears to be constitutively active in T-REx293 cells but not in BMMs. After 1 h without substrates or bicarbonate, pyruvate and hydroxybutyrate inhibit constitutive OMDE, as expected for a shift of CoA from long-chain acyl-CoAs to other CoA metabolites. In the presence of bicarbonate, pyruvate strongly enhances OMDE, which is then blocked by β-hydroxybutyrate, bromopalmitate/albumin complexes, cyclosporines, or edaravone. After pyruvate responses, T-REx293 cells grow normally with no evidence for apoptosis. Fatty acid-free albumin (15 μM) inhibits basal OMDE in T-REx293 cells, as do cyclosporines, carnitine, and RhoA blockade. Surprisingly, OMDE in the absence of substrates and bicarbonate is not inhibited by siRNA knockdown of the acyltransferases, DHHC5 or DHHC2, which are required for activated OMDE in patch clamp experiments. We verify biochemically that small CoA metabolites decrease long-chain acyl-CoAs. We verify also that palmitoylations of many PM-associated proteins decrease and increase when OMDE is inhibited and stimulated, respectively, by different metabolites. STED microscopy reveals that vesicles formed during constitutive OMDE in T-REX293 cells have 90 to 130 nm diameters. In summary, OMDE is likely a major G-protein-dependent endocytic mechanism that can be constitutively active in some cell types, albeit not BMMs. OMDE depends on different DHHC acyltransferases in different circumstances and can be limited by local supplies of fatty acids, CoA, and long-chain acyl-CoAs.

## Introduction

Eukaryotic cells use endocytosis (1, 2) to carry out a wide range of cellular functions. Phagocytosis and macropinocytosis are used to internalize large particles and fluid volumes (3). Micro-pinocytotic mechanisms generate vesicles from the plasma membrane (PM) with diameters of 40 to 200 nm (4), thereby internalizing integral PM proteins, PM-associated proteins, and various other types of particles, as well as fluid (5). In doing so, micro-pinocytotic mechanisms re-localize membrane and membrane proteins from the cell surface into internal membrane systems from which membrane and protein can be recycled, degraded, and/or used for cell signaling functions. Innumerable studies show that micro-pinocytosis can occur by diverse mechanisms, ranging from the coordinated function of clathrin, dynamins, and other adapter proteins to the lateral reorganization of plasma membrane lipids and proteins into domains, often with additional involvement of actin cytoskeleton to direct budding and vesiculation (6). Dynamins can drive some forms of endocytosis without clathrin (7). However, one such example has recently been questioned. In contrast to a long-time assumption of the field, it was recently described that dynamin-2 may not support the internalization of caveolae, but rather restrain caveolae internalization (8). Another instructive example concerns turnover of PM in two cell types used routinely in cell biological studies. For HeLa cells, it has been proposed that nearly all PM turnover under standard cell culture conditions occurs via clathrin/dynamin-dependent endocytosis (9). In mouse fibroblasts, by contrast, the deletion of all three dynamins results in no clear change of fluid phase endocytosis (10), and we verify this in Supplemental Fig. S1. Importantly, analysis of fluid phase dextran uptake by confocal optical methods can readily detect clathrin/dynamin endocytosis (e.g.(11)). Thus, it remains enigmatic whether (i) different endocytic mechanisms operate differently in these different cell types, (ii) different endocytic mechanisms compensate one another when one is manipulated by genetic means, or whether (iii) clathrin/dynamin endocytosis is only a small fraction of total PM turnover in fibroblasts.

Using patch clamp to monitor PM area changes and to manipulate the contents of the cytoplasm, we have been impressed by the large magnitudes and speeds with which endocytic responses can occur via clathrin/dynamin-independent mechanisms (12). We have documented in multiple cell types the occurrence of massive endocytosis that internalizes more than 50% of the PM into relatively small vesicles (i.e. less than 240 nm) within one to two minutes (12). We have described these responses in primarily three circumstances—in the wake of large cytoplasmic Ca elevations, during activation of G-proteins by nonhydrolyzable GTP analogs, and with activation of oxidative metabolism in mitochondria by substrates such as succinate. In all cases, we have verified that the membrane that internalizes binds amphipathic compounds less well than membrane that remains at the cell surface (13, 14). Given that more ordered membrane domains internalize, we suggest that this form of endocytosis may productively be called <u>o</u>rdered <u>m</u>embrane <u>d</u>omain <u>e</u>ndocytosis (OMDE). Potentially, this term can be used in the future to include the various forms of endocytosis otherwise dubbed ‘raft’ (15), ‘CLIC’ (16) and ‘bulk’ (17) endocytosis. All of these endocytic mechanisms are dynamin/clathrin-independent and may have dependence on actin cytoskeleton in addition to membrane lipid and protein ordering.

While patch clamp results suggest that OMDE can be remarkably powerful when activated, available data does not establish that OMDE is active constitutively in cells or under standard cell culture conditions. Rather, work to date indicates that OMDE becomes active when cellular conditions promote palmitoylation of membrane proteins that subsequently can coalesce into ordered domains. As already noted, mitochondrial oxidative stress appears to promote this process, and our working hypotheses about the underlying mechanisms are outlined in cartoon form in Fig. 1. In brief, mitochondria accumulate the ubiquitous enzymatic cofactor, coenzyme A (CoA), to high concentrations with respect to the cytoplasm (18, 19). As mitochondria generate reactive oxygen species (ROS) in response to enhanced cytochrome oxidase activities and Ca accumulation, transient openings of permeability transition pores (PTPs) allow CoA to escape to the cytoplasm, promoting generation of long-chain acyl-CoAs. In dependence on acyl-CoA accumulation, as well as other signaling mechanisms that are only partially elucidated, key PM proteins are evidently rapidly palmitoylated to promote OMDE.

**Figure 1.**
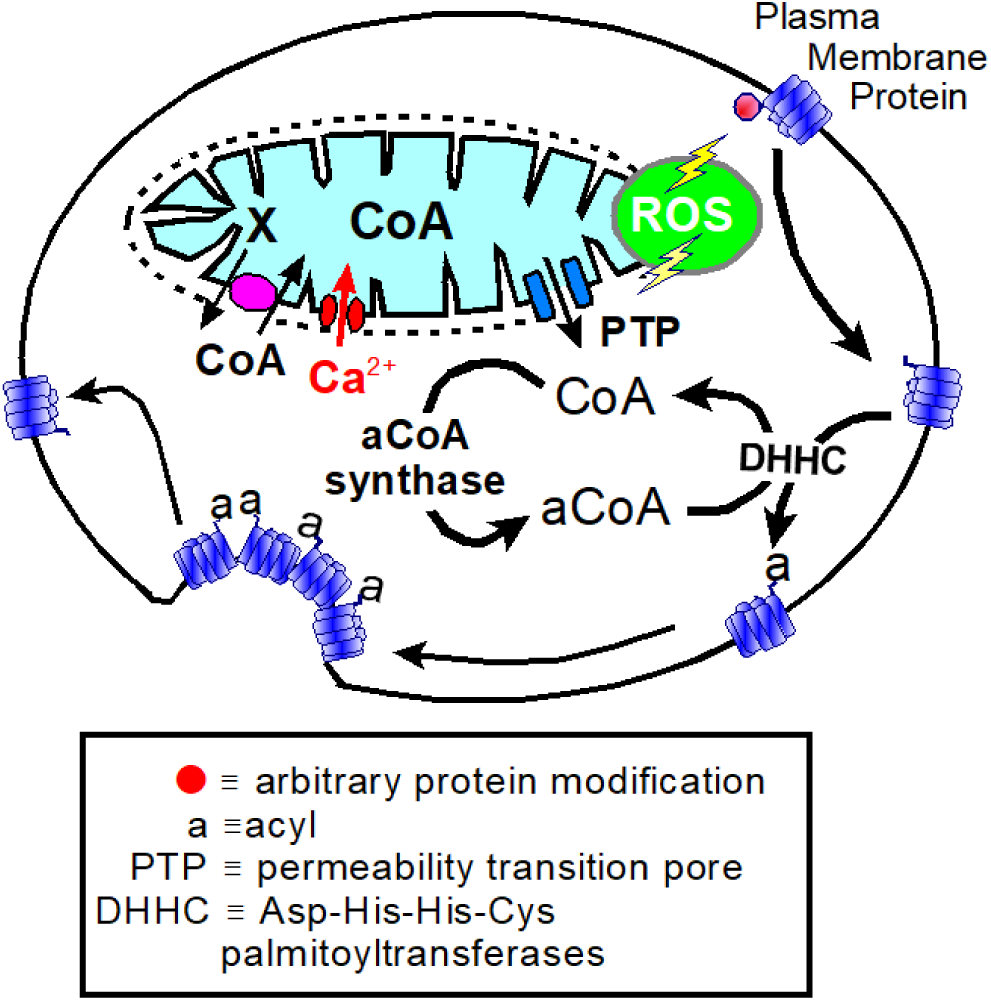
Working hypotheses. Mitochondria accumulate coenzyme A to a high concentration with respect to cytoplasm and release it into the cytoplasm during metabolic stress, e.g. post-ischemic stress (21, 88), but also via constitutive PTP openings Cytoplasmic CoA (89) and acetyl-Co A (90) concentrations fluctuate enough to influence enzymes that use them as cofactors and substrates. Thus, both molecules have second messenger functions. CoA is released from mitochondria via transient PTP openings that occur physiologically as mitochondria accumulate Ca and oxidize critical cysteines (e.g. (91)). Elevations of cytoplasmic CoA promote acyl-CoA synthesis and subsequently DHHC acyltransferase activity. DHHC-mediated palmitoylations are regulated by small G-proteins that modify membrane cytoskeleton function, ROS signaling, phosphorylations (92), and conformational changes of the PM and PM proteins. Reversible palmitoylations of PM proteins can promote their coalescence into ordered membrane domains, commonly called ‘rafts’, thereby supporting the budding of ordered domains inwardly to eventually form endocytic vesicles (93). Palmitoylation-dependent ordered membrane domain endocytosis (OMDE) is long known to mediate uptake of proteins and particles by cells. Examples include the uptake of hyaluronic acid by CD44 (83), uptake of oxidized LDL and fatty acids by CD36 (84), uptake of nanoparticles and proteins by heparan sulfate proteoglycan receptors (94, 95), and uptake of multiple viruses (96). Additionally, the clustering of some membrane transporters, independent of ligands, promotes their endocytosis in a cargo-dependent manner without involvement of classical endocytic proteins (32, 97).

To address how and when this occurs in cultured cells without patch clamp, we describe here studies of fluid phase endocytosis of small fluorescent dextrans (3-10 kD) in which we have tested acute effects of factors that promote or hinder OMDE in patch clamp experiments. Further to this end, we compare the function of endocytosis in two very different cell types, T-REx293 cells and differentiated murine bone marrow macrophages (BMMs). To develop clear perspectives on OMDE, we describe first via patch clamp that OMDE activated by oxidative metabolism in BMMs is more powerful than in other cells, internalizing 75% of the PM in 2 min and requiring expression of the acyltransferase, DHHC5. While less powerful, we show that OMDE activated by GTPγS in T-REx293 cells is highly dependent on actin cytoskeleton when membrane tension is high, but not when membrane tension is relieved. Then, analyzing fluid phase endocytosis of dextrans unperturbed cell cultures, we show that constitutive (basal) endocytosis in T-REx293 cells is significantly activated by palmitate/albumin complexes and reduced by inhibitors of palmitoylation, as well as by RhoA G-protein inhibition and cyclosporines that inhibit mitochondrial PTPs. We go on to show that constitutive endocytosis in T-REx293 cells can be strongly activated by extracellular pyruvate in the presence of bicarbonate. OMDE is inhibited by low concentrations of fatty acid-free albumin (15 μM) and can be restored by albumin/palmitate complexes. While pyruvate is stimulatory, some mitochondrial substrates are inhibitory. Finally, we show via STED microscopy that vesicles formed in OMDE are 90 to 120 nm in diameter. All results for T-REx293 cells, but not for BMMs, are consistent with mitochondria-dependent OMDE constituting a large fraction of basal PM turnover. However, the constitutive OMDE activity appears to use DHHC acyltransferases besides DHHC5 and DHHC2. We conclude that constitutive (basal) PM turnover in these different cell types occurs by different mechanisms and that OMDE in BMMs must be activated by cell signaling mechanisms that are not constitutively active.

## Materials and Methods

### Mice

The UT Southwestern Medical Center Animal Care and Use Committee approved all animal studies. The DHHC5 gene-trapped mice were from the same line used previously in which expression of DHHC5 in all tissues tested was reduced by >80% (20, 21). Equal numbers of male and female mice were employed in experiments with no significant differences in results.

### Primary Cells, Cell Culture and Transfection

#### Isolation of Lung Fibroblasts

Lung fibroblasts of DHHC5-GT animals and their WT controls were isolated as described elsewhere (22). In brief, animals were euthanized, sterilized with ethanol, and transferred to the laminar flow. Lungs were excised, cut in 1mmx1mm pieces, and placed in a grated Petri-dish. 10 mL of high-glucose DMEM supplemented with 10% heat-inactivated FBS, 50U/mL Penicillin/ and 50µg/mL streptomycin were added carefully, and dishes were placed in a humidified incubator with 5% CO_2_ at 37°C. After 7-10 days, tissue pieces were removed, and outgrown cells were subjected to experiments after they reached 90% confluency.

#### Isolation and Differentiation of Bone Marrow Macrophages

Bone marrow cells of DHHC5-GT animals and their controls were isolated as described (23). In brief, animals were euthanized, sterilized with ethanol, and transferred to the laminar flow. Hind limbs were removed, and the femur and tibia were cleaned rigorously. Bone marrow was flushed out of the bones with 1x PBS and was collected in conical tubes on ice. After resuspension of bone marrow pieces, cells were filtered through a 75 µm cell strainer and the filtered suspension was centrifuged for 5 min at 500xg. The cell number was then determined using a hematocytometer. Cells were seeded at a density of 1×10^6^/six-well well in high-glucose DMEM supplemented with 10% heat-inactivated FBS, 50U/mL penicillin/ and 50µg/mL streptomycin and 25 ng/mL M-CSF (PreproTech, Cranbury, NJ). Cells were employed in experiments after 7 days. For patch clamp experiments, cells were trypsinized and employed without reattaching to dishes.

#### Isolation of Peritoneal Macrophages

Peritoneal macrophages of DHHC5-GT mice and their WT controls were isolated as described previously (24). In brief, animals were euthanized, sterilized with ethanol and the skin layer was carefully removed from the abdomen. The inner skin lining of the peritoneal cavity was then carefully punctured and 5 mL of PBS supplemented with 3% FBS were injected. The peritoneal cavity was massaged for about 30 s, the isolation medium was re-aspirated, and cells were transferred to 50mL of solution in a conical tube. The tube was centrifuged for 5 min at 500xg. Cells were resuspended in DMEM supplemented with 10% FBS and thereafter employed in electrophysiological measurements.

#### Cell Culture and Transfection

T-REx 293 cells were grown in high-glucose DMEM supplemented with 10% heat-inactivated FBS, 50 U/mL penicillin/ and 50µg/mL streptomycin in a humidified incubator with 5% CO_2_ at 37°C. Transfection of T-REx293 cells with siRNA was performed using Lipofectamine RNAiMAX (Thermo Fisher Scientific, Waltham, MA) according to the manufacturer’s instructions. siRNAs targeting DHHC 2 and 5 were obtained from Thermo Fisher Scientific (Waltham, MA) and cells were subjected to experiments 48 h after transfection.

#### Generation of DHHC5/2 knock-out cells

CRISPR knock-out cell lines were produced using established methods (19, 20). The T-REx293 cell line was transiently transfected with LentiCRISPRv2 (Addgene, #52961). The sgRNA sequences used for cloning LentiCRISPRv2-DHHC5 were: DHHC5-forward primer, CCGTGAATAACTGTATTGGTCGC and DHHC5-reverse primer, AAACGCGACCAATACA GTTATTCAC. The sgRNA sequences used for cloning LentiCRISPRv2-DHHC2 were: DHHC2-forward primer, CACCGCGTAGTAGGACCAGCCGAGC and DHHC2-reverse primer, AAACGCTCGGCTGGTCCTACTACGC. The sequencing primer of LentiCRISPRv2 was hU6F: GAGGGCCTATTTCCCATGATT, which was used to verify the sgRNA/Cas9-mediated DHHC2 or DHHC5 gene knockout clones. After the appropriate plasmids were transfected, cells were cultured with medium containing puromycin (10 µg/ml) for one week to select for the surviving cells with the desired plasmids for knockdown of the two genes. For dual knockdown of DHHC2 and DHHC5, cells were co-transfected with LentiCRISPRv2-DHHC5 and LentiCRISPRv2-DHHC2, followed by the relevant selection procedures.

### Patch clamp, and capacitance measurement

Axopatch 1C patch clamp amplifiers were employed using our own software for capacitance and conductance recording via square-wave voltage perturbation (25). Unless stated otherwise, solution compositions were the same as employed previously (21). All patch clamp experiments were performed at 0 mV and 35°C. Extracellular solution changes were performed by abruptly moving the microscope stage so as to place the patch-clamped cell directly in front of 1 of 4 square pipettes (1 mm) with solution steams maintained by gravity at velocities of 2 to 5 cm/s. In all experiments in which two or more protocols were compared using a single batch of cells, protocols were performed in a randomized fashion.

### STED microscopy

STED microscopy was performed in the UTSouthwestern O’Donnell Brain Institute NeuroMicroscopy Core Facility using an Abberior Instruments STED microscope. The Abberior Facility Line system is a 3-D STED super-resolution imaging system. Super-resolution images were acquired from fixed samples, prepared as described. 30 to 40 nm isotropic resolution was verified in our samples. Adaptive Illumination (RESCue, DyMIN and MINFIELD) was employed with a 60X (NA1.42 WD 0.15mm) Oil Objective. Pulsed 485 nm, 561 nm, and 640 nm lasers were employed. 595 nm and 775 nm STED depletion lasers were employed.

### Immunoblotting

For protein isolations cells were washed 3x with pre-cooled PBS and homogenized in pre-cooled RIPA buffer (containing in mM: 150 NaCl, 50 Tris–HCl [pH 8.0], 5 EDTA, 1 EGTA; Triton X-100 1% [vol/vol], deoxycholate 0.5% [wt/vol], SDS 0.1% [wt/vol], and protease inhibitor cocktail from Roche) and lysed for 30 minutes at 4°C. Lysates were cleared at 20,000xg for 15 minutes and subjected to SDS PAGE and subsequent immunoblotting (26). Anti-sera: Anti-ZDHHC5 antibody, produced in rabbit, HPA014670, was from Sigma. Anti-ZDHHC2 antibody, produced in mouse, Clone 1035A (27), was a gift of Dr. Masaki Fukata, Okazaki. Since the DHHC2 antibody provided by Dr. Fukata is no longer available, nor is a comparable antibody available, we subsequently evaluated knockdown of DHHC5 and DHHC2 by qPCR, as described below. Anti-actin was purchased from Santa Cruz (Dallas, TX).

### RNA Isolation and Real-Time PCR

RNA isolation and Real-Time PCR were performed as described (28). In brief, total RNA was isolated using Qiagen RNeasy Plus Mini Kits according to manufacturer’s instructions and 1 µg of RNA was reverse transcribed employing the High-Capacity cDNA Reverse Transcription Kit. qPCR was performed on a BioRad (Hercules, CA) CFX96 real time system. Primers for DHHC2 and DHHC5 RNA were as follows: DHHC2 forward primer, TCCCGGTGGTGTTCATCAC. DHHC2-reverse primer, CAACTTGTTCGCCAGTGTTTTC (product length: 99bp). DHHC2-forward primer 2, AACACTGGCGAACAAGTTGTG. DHHC2-reverse primer 2: AGATGGGAAGATCCTTGGCTG (product length: 211bp). DHHC5 forward primer 1, CACCTGCCGCTTTTACCGT. DHHC5-reverse primer 1, CGGCGACCAATACAGTTATTCAC (product length: 111bp). DHHC5-forward primer 2, AACTGTATTGGTCGCCGGAAC. DHHC5 reverse primer 2, AACACACCCATAATGTGGGCT. Knockdowns amounted to 70 and 72% for DHHC2 and DHHC5, respectively.

### Acyl-RAC Procedure for Mass Spectrometry

The Acyl-RAC procedure was conducted as described elsewhere (25, 26). In brief, T-REx293 cells were grown to 90% confluency in 10 cm dishes and then subjected to the indicated treatments. The cells were washed and lysed in 2 mL of lysis buffer (containing in mM: 50 Tris pH 7.2, 5 EDTA, 150 NaCl, 10 NEM, Protease Inhibitor Cocktail (Roche, Basel, Switzerland), 2 PMSF) using Dounce homogenization. Lysates were then centrifuged at 200,000xg for 30 minutes to separate the membrane and soluble fractions.

Membrane fractions were resuspended in lysis buffer supplemented with 1.7% Triton-X-100 using a 23 G needle. Solutions were then incubated under rotation at 4°C for 1 hour and collected by centrifugation at 250xg for 5 minutes. For acetone precipitation, 3 volumes of ice-cold acetone were added, and the solution was incubated for 30 minutes at −20°C. The solution was then centrifuged at 3000xg for 5 minutes to collect the pellet. The pellets were washed 3 times with pre-cooled 95% ethanol and dried for 5 minutes. The pellet was resuspended in the same volume as before the acetone precipitation in blocking buffer (containing in mM: 20 HEPES pH 7.7, 5 TCEP, 5% [wt/vol] SDS) and subjected to BCA. After adjusting the protein concentratio to 2 mg/m, extracts were incubated at 30°C for 30 minutes, 10 mM NEM were added, and extracts were incubated rotating for 1 hour at 37°C. For acetone precipitation, 3 volumes of ice-cold acetone were added, and the solution was incubated overnight at −20°C. The solution was then centrifuged at 3000xg for 5 minutes to collect the pellet. The pellets were washed 3 times with pre-cooled 95% ethanol and dried for 5 minutes. Pellets were resuspended in column buffer (containing in mM: 20 HEPES at pH 7.2, 1 EDTA, 1% [wt/vol] SDS, and the tinal protein concentration was adjusted to 2 mg/mL.

Samples were split in two, and 300 µL of thiol-sepharose slurry washed with column buffer was added to each sample. Corresponding fractions were either treated with a final concentration of 0.7 M freshly prepared hydroxylamine (pH 7.2) or with 0.7 M Tris (pH 7.2). Samples were incubated under rotation overnight (23°C). Beads were allowed to settle by gravity, supernatant was collected, and beads were washed 3 times with 1xPBS. The last wash was removed, 150 µL of 2x TEABC buffer 1 (containing in mM: 50 TEABC) and 25 µg of trypsin/sample were added, and samples were incubated under rotation at 37°C overnight. Beads were allowed to settle by gravity, supernatant was collected, and beads were washed 3 times with 1xPBS. To elute proteins, 150 µL of 2x TEA-BC buffer 2 was added (50 mM TEABC, 10 mM TCEP), followed by incubation at 37°C for 30 min and collection of supernatants for downstream applications.

### Fluid phase dextran uptake assay

T-REx 293 cells were seeded on Matek glass bottom dishes at least 48 h prior to experiments, such that cell confluency was 80-90% at the time of experiments. BMMs were trypsinized on day 6 of differentiation and seeded onto Matek glass bottom dishes for an additional 24 - 48 h. The cells were then washed and pre-treated for 30 minutes with the saline solution described. For serum/substrate-free experiments, cells were washed 3x in a saline solution containing in mM: 140 NaCl, 5 KCl, 10 HEPES, 0.6 MgCl_2_, 2 CaCl_2_, and 0.6 NaHPO_4_, at pH 7.2. Cells were then incubated for 30 min or 1 h, as indicated, with the same saline solution followed by a chosen treatment, as described in Results, before application of the same solution with 100 μM TexasRed lysine-fixable dextran 3000 MW (Thermo Fisher Scientific, Waltham, MA). Uptake times of either 20 or 4 min were employed, as indicated in Results. Thereafter, cells were rapidly placed on ice and rinsed twice with cold solution. In some experiments, cells were stained with WGA 488 for 5 min to visualize the PM, followed by three cold washes. All cells were fixed for 10 minutes with 4% PFA, washed with PBS, stained with Hoechst dye according to the manufacturer’s specifications and then overlayed with CitiFluor Mountant AF1 Solution (Electron Microscopy Sciences, Hatfield, PA).

### Fluorescence imaging

Imaging at near confocal resolution was carried out with an Aurox Clarity laser-free microscopy system (Abingdon, Oxfordshire, OX14 3DB, UK) (19), employing a Nikon Eclipse TE 2000-S inverted microscope with an APO LWD 40x 1.15 NA water immersion lens. The 50% point spread was 0.7 and 1.6 microns in the lateral and vertical directions, respectively, as determined with a TetraSpeck Fluorescent Microspheres Size Kit (Invitrogen). This compares to 0.6 and 2.9 microns for a 40x 0.95 NA air lens employed on a standard spinning disc microscope (Nikon CSU-W1 spinning disk with Hamamatsu Orca Fusion camera, Nikon CSU-W1 SoRa: Quantitative Light Microscopy Core Facility, UTSouthwestern). Equivalent images obtained from the same slide (FluoCells prepared slide #3, Invitrogen) with these two imaging systems were provided previously (29).

### Analysis of dextran uptake assays

All bar graphs depicting dextran uptake correspond to the analysis of 12 or more fields of view from at least two dishes of cells fixed as described. Typically 4 images were taken from each of the dishes to be compared, and the procedure was repeated 3 times in the same sequence. Fig. 2 illustrates our routine analysis of the dextran uptake assays using a relatively complex sample in which cell density reached confluence in some regions and was sparse in others. Figs.2A to 1C show the fluorescent dextran and Hoechst dye images of a field of T-REx293 cells (∼150 cells) that was treated and fixed as described above. Using ImageJ software, a threshold fluorescence was set high enough to eliminate background fluorescence in cells fixed without dextran (not shown). Particles were then identified and outlined, as shown in Fig. 2C using an upper limit set to 2 square microns. Particles were counted in this without setting a lower particle size limit, the influence of which is described subsequently. The number of nuclei was determined as described in Figs. 2C and 1D. As apparent in the Hoechst image in Fig. 2C, nuclei often overlapped in regions where the cell culture was nearly confluent. As shown in Fig. 2C, the ImageJ software was then unable to accurately separate nucleus images, so that automated nucleus counting became impossible. Therefore, we used two other strategies to determine the number of nuclei. In some cases, we counted nuclei manually by marking them individually via ImageJ. Second, we determined the total nuclear area, as defined by nuclei outlines, as in Fig. 2D. We then divided the total area by the average nucleus area determined from 20 to 40 individual nucleus area measurements. As indicated below Fig. 2D, the two approaches yielded values that did not differ by more than 5 to 6%, 153 versus 144 nuclei in this case.

**Figure 2.**
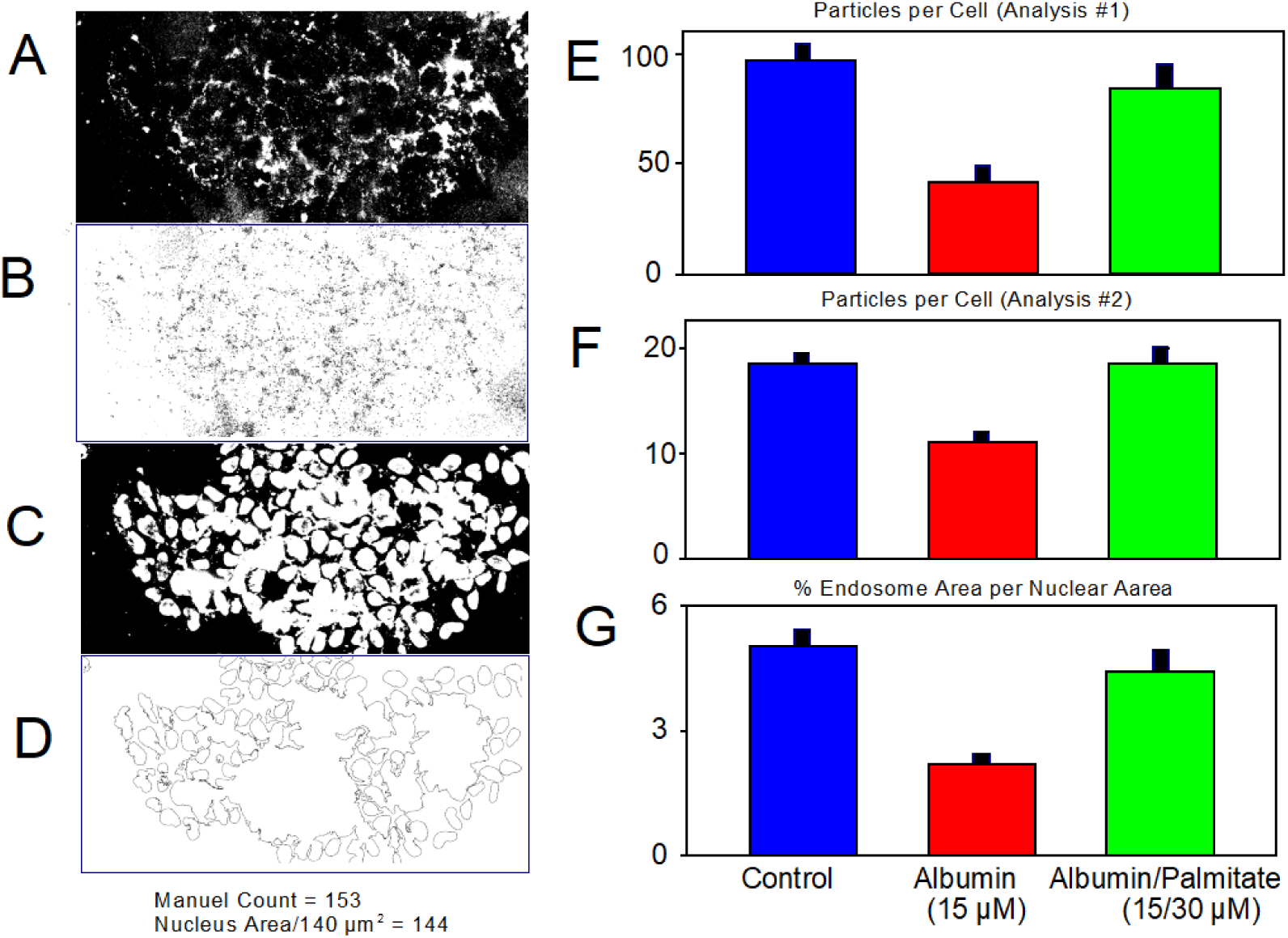
Analysis of 3kD TexasRed^TM^-dextran uptake into T-REx293 cells for 4 min. To determine cell numbers, cells were labelled with Hoechst 33343 dye after terminating uptake with cold solution. We illustrate our ImageJ-based analysis with an example in which cells are distributed unevenly and partially very densely. **A.** Fluroescence image of dextran uptake with a threshold set to exclude autofluorescence. **B.** Outlines of all punctae rendered by ImageJ up to a maximal particle size of 2 μm^2^. Since the lower threshold excludes light from puncta edges, puncta dimensions become compressed. Therefore, we do not enforce a lower limit on puncta sizes. **C.** Image of Hoechst-stained nuclei for the same field. A manual count of nuclei yielded a value of 153. **D.** Image of nuclei borders determined by ImageJ. Typical for high cell density, the outlines of neighboring nuclei merge so that automated nuclei counts are inaccurate. Therefore, we determined the total nuclear area from outlines and divided by an average nucleus area, determined from >50 measurements. The estimate was 144 nuclei. Similar to 5 other cases, differences between this automated cell count and manual counts was less than 15%. **E-G.** Representative analysis of dextran uptake. The results are from Fig.7, showing the effect of FA-free albumin (15 μM) and palmitate-loaded albumin (15/30 μM) on dextran uptake. 7. **E**. Particle counts per cell, determined as in in panels A-D. Average particle counts for 4 min dextran uptake were ∼100 particles per cell. Counts were reduced by 63% with albumin exposure, while they were only marginally decreased by albumin/palmitate complexes. **F.** Particle counts with minimum particle diameter set to 0.2 microns. Control particle counts are reduced to about 20 per cell when the size limit is set to the wavelength of light. As described in Fig. 11, analysis with STED microscopy verifies that particle counts without a lower dimension limit are more correct. We note also that the imposition of a lower particle size limit artificially distorts the magnitude of the inhibitory effect of albumin (see Panel F versus Panel E). **G.** Representative for more than 20 observations, changes of dye uptake, as accessed by particle counts, are essentially the same when dye uptake is quantified as endosome (particle) area divided by total nuclear area.

Figures 2E to 2G illustrate the analysis of dextran uptake using different criteria. The data shown are from the experiment described subsequently in Fig. 8B, demonstrating that dextran uptake is markedly inhibited by fatty acid free albumin (15 μM) but not by 2:1 palmitate/albumin complexes at the same concentration. Figs. 2E and 2F show results quantified as particles per cell for the three conditions, both without setting a lower particle size threshold and with a lower particle size limit set to 0.25 square microns (i.e. a diameter of 500 nm). In the latter case, the number of apparent particles is 5-times less and the apparent dextran uptake is reduced by only 40% versus 60%. Based on our subsequent analysis using STED microscopy, we concluded empirically that the former analysis (Fig. 2E), which generates higher particle numbers, is in fact more accurate. This can be justified by the fact that the thresholding procedure employed results distorts the dimensions of small particles so as to be smaller than the wavelength of light. Finally, Fig. 2G shows the same experimental results quantified as total particle area per total nuclear area, instead of particles per cell. The results are quantitatively very similar to results for particle numbers, indicating simply that average particle size does not change with FA-free albumin treatment.

### Chemicals

Unless stated otherwise, chemicals were from Sigma-Millipore (St. Louis, MO) and were the highest grade available.

### Experimental Design and Statistics

All patch clamp results presented were similar in sets of experiments performed on at least 3 different days. When practical, experiments were performed in a blinded or double-blinded fashion. In each uptake assay, at least two dishes of cells were analyzed for each experimental condition, and similar results were obtained in at least two different sets of experiments. As already noted, all bar graphs depicting dextran uptake include at least 12 images from two dishes of cells. Significance was determined via Students’ t-test, except in rare cases when results were not normally distributed. In those cases, significance was determined by Mann-Whitney Rank Sum test. Error bars represent Standard Error.

We considered and tested several statistical methods to analyze acyl-RAC results using mass spectrometry to quantify proteins. Results presented in Fig. 9 are all from triplicate cell samples prepared in parallel under identical conditions. Given that mass spectrometry results can be relatively variable from sample to sample, we employed the median value from the three measurements in subsequent calculations. From these measurements as well as quantification of Western blots in parallel, we found that caveolin-1 was consistently highly enriched in samples and was never observed to change significantly with treatments carried out. This outcome was not unexpected, given that caveolins are palmitoylated at minimally three cysteines (30) and that palmitoylation at a single cysteine results in pull-down in the acyl-RAC procedure employed. Therefore, we normalized protein in each set of measurements to the amount of caveolin-1 present in the sample. Results were qualitatively very similar, but more prone to variability, when quantified from the absolute protein counts.

## Results

### DHHC5/2-dependent OMDE: Activation by GTPγS and oxidative metabolism in patch clamped T-REx293 cells and BMMs

Figure 3 describes large-scale OMDE that can be activated without Ca elevations in T-REx293 cells and BMMs. Fig. 3A presents representative capacitance and conductance records from a T-REx293 cell that was opened via patch clamp with a pipette solution containing GTPγS (0.2 mM), MgATP (10 mM) and EGTA (10 mM) with no added Ca. Free cytoplasmic Ca is thus essentially zero. Other solution constituents were as in previous experiments (31) and were not critical for OMDE function. After opening the cell by suction, the cell was moved quickly into a solution stream that was warmed to 35°C. Without discernable changes of cell conductance, membrane capacitance (i.e. PM area) decreases by about 40% over 8 min. As described previously (32), this occurs more than 5-fold faster, and can be enhanced, when T-REx293 cells overexpress the cardiac NCX1 Na/Ca exchanger, which is palmitoylated at one cysteine close to the PM (32).

**Figure 3.**
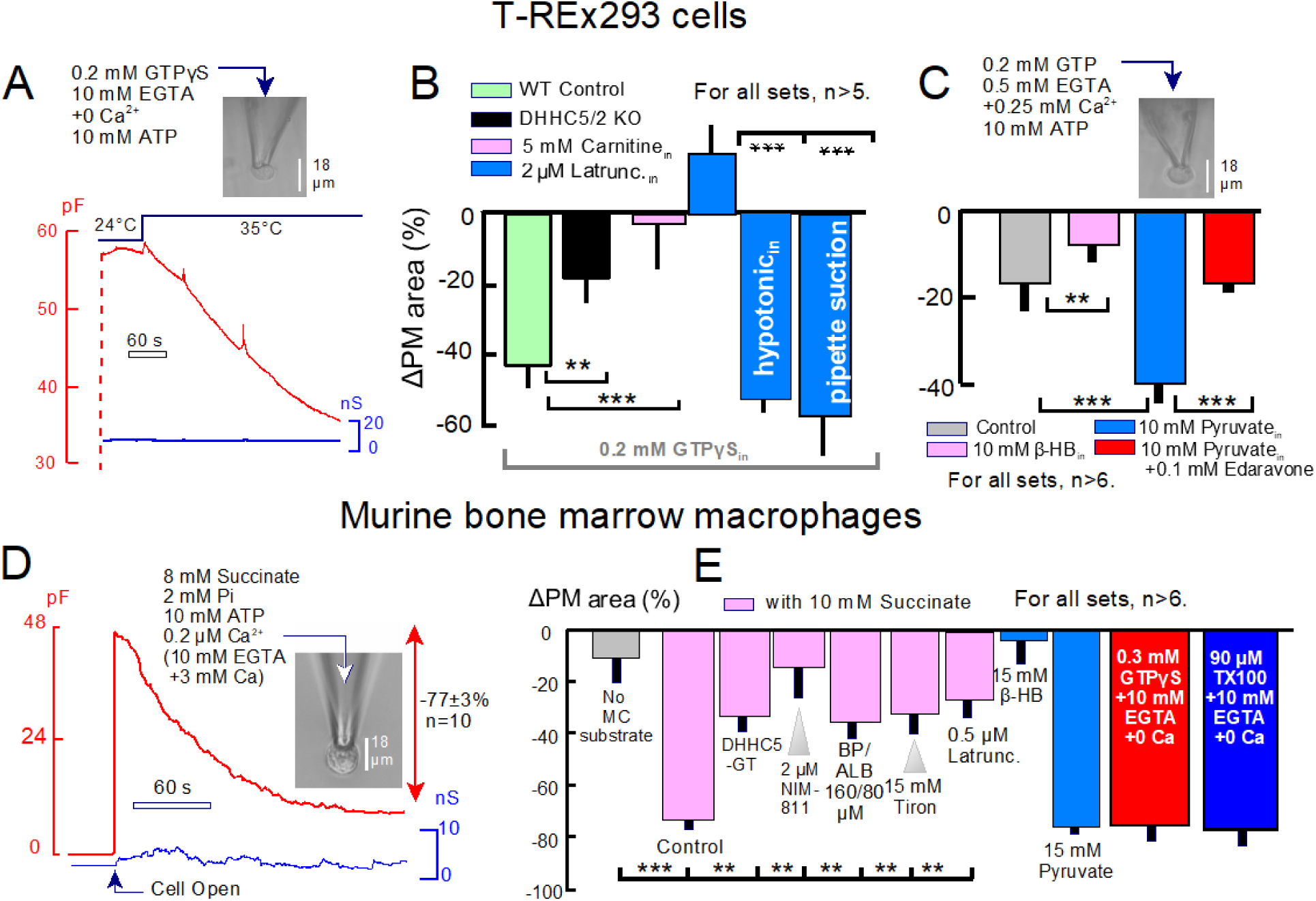
Patch clamp analysis of OMDE activated by cytoplasmic GTPγS and by mitochondrial substrates (succinate or pyruvate) in T-REx293 cells (A-C) and in cultured murine BMMs (D-E). **A.** Typical recording of T-REx293 cell capacitance (red) and conductance (blue) after cell rupture in the presence of GTPγS (2 mM) in the pipette solution with 10 mM EGTA, no added Ca, and 10 mM Mg ATP. Upon moving the cell into a warm solution stream (35°C), cell capacitance (i.e. PM area) begins to fall within seconds and decreases by 40% over 8 min. Cell conductance remains unchanged. **B.** Composite results for experiments employing the protocol of Panel A. From left to right, average PM loss amounts to 42 % (green), PM loss is decreased from 40% to 18% in a cell line in which the acyltransferases DHHC5 and DHHC2 were deleted by CRSPR (black), PM loss is decreased by 90% when 5 mM carnitine (5 mM) is included in the pipette (pink), and PM loss is not only blocked, but replaced by a small PM expansion response, when latrunculin (2 μM) is included in the pipette (blue). The effect of latrunculin is completely reversed by promoting cell shrinkage with a 30% hypotonic cytoplasmic solution or by applying enough negative pressure to the pipette to induce a visible cell shrinkage (blue). **C.** PM loss after T-REx293 cell opening with a pipette solution containing GTP (0.2 mM), EGTA (0.5 mM) with Ca (0.25 mM, free Ca ≈ 0.3 μM), and MgATP (10 mM). PM loss over 5 min amounts to 18% in control cells. Loss is reduced to 8 % in cells perfused with β-HB (10 mM), and loss is increased to 40% in cells perfused with 10 mM - pyruvate. Inclusion of antioxidant, edaravone (0.1 mM), with pyruvate completely suppresses the pyruvate-dependent endocytosis. **D.** Representative record of BMM cell capacitance (red) and conductance (blue) changes upon cell rupture, just after the cell was warmed to 35°C. With a pipette solution containing succinate (8 mM), Pi (2 mM), Mg ATP (10 mM) and free Ca clamped to ∼0.2 μM (10 mM EGTA with 3 mM Ca), BMM capacitance amounts initially to 48 pF. Capacitance declines over 4 min to ∼10 pF while conductance remains stable. **E.** From left to right, loss of PM area amounts to only 10% when succinate is not included in the pipette (gray) versus 77% with succinate (pink). PM loss is decreased by 65% in BMMs from animals with gene-trapped DHHC5 acyltransferase (pink), PM loss is decreased 77% when the mitochondria-specific cyclosporin, NIM-811 (2 μM) is included in solutions on both membrane sides (pink). PM loss is decreased by 60% when cells are incubated with brompalmitate/ albumin complexes for 10 min prior to recording (160/80 μM, pink), decreased by 63% when the antioxidant, tiron (15 mM) is included in the pipette (pink), and decreased 70% when cells are pretreated with latrunculin (0.5 μM) for 10 min (pink). PM loss was less than 10% when the pipette solution contained β-hydroxybutyrate (15 mM), instead of succinate (blue), while pyruvate (15 mM) promoted an equal PM loss to that occurring with succinate. PM loss was similarly large when GTPγS was used to induce OMDE in the absence of mitochondrial substrates. Finally, PM loss was similarly large when rapid application of tritonX-100 (90 μM) was used to induce massive endocytosis by promoting PM phase separations (12).

Figure 3B illustrates additional key features of OMDE activated by GTPγS that were not described previously. From left to right, the responses in cells without NCX1 expression amounted on average to a 40% loss of PM over 8 min. Using T-TEx293 cells in which two DHHCs known to reach the PM, DHHC5 and DHHC2 (33–35), are knocked out by CSPR (see Supplemental Fig. S2), the PM loss is decreased to 18%. Cytoplasmic carnitine is a substrate of carnitine palmitoyl transferase, which converts long-chain acyl-CoAs to acyl-carnitines that are transported into mitochondria and resynthesized to acyl-CoAs (36). The endocytic response is decreased to less than 5% when 5 mM carnitine is included in the pipette solution, presumably by depleting the cytoplasmic pool of long-chain acyl-CoAs. This result supports our suggestion that long-chain acyl-CoAs can limiting for the endocytic responses under the conditions of these experiments (37). When actin cytoskeleton is disrupted by treating cells for 10 min with latrunculin (0.5 μM), and including latrunculin (2 μM) in the pipette, the PM loss is blocked and replaced by a PM expansion response of about 25%. Nevertheless, the loss of PM induced by GTPγS can be fully restored by performing the same experiments with a 30% hypotonic solution to induce cell shrinkage (5^th^ bar graph). Also, the responses are restored simply applying enough suction to the pipette tip to cause a detectable amount of cell shrinkage (6^th^ bar graph). From these results, we suggest that latrunculin may inhibit endocytosis by allowing the PM to expand laterally and stretch, thereby generating lateral membrane tension that hinders membrane budding. This interpretation was proposed already decades ago to explain actin engagement in clathrin-mediated endoctyosis (38). Importantly, these results indicate that G-proteins activated by GTPγS promote endocytosis by mechanisms besides the modulation of membrane actin cytoskeleton.

Figure 3C illustrates two manipulations of mitochondrial metabolism that are central to our analysis of OMDE in T-REx293 cells in culture. When cells are opened in the presence of GTP (0.3 mM), MgATP (10 mM) and minimal Ca buffering (0.5 mM EGTA with 0.25 mM Ca, giving about 0.25 μM free Ca), PM loss amounts to about 17% over 8 min. This loss is decreased by 60% when the mitochondrial substrate, β-hydroxybutyrate (β-HB, 10 mM) is included in the pipette (2^nd^ bar graph), but the loss is enhanced 2.5-fold when the same concentration of the mitochondrial substrate, pyruvate, is included in the pipette (3^rd^ bar graph). Thus, the metabolism of mitochondrial substrates is a major determinant of OMDE, and the extent to which ROS are formed during their metabolism is one potentially important determinant of OMDE progression. As shown in the 4^th^ bar graph, the hydrophobic antioxidant, edaravone, also called MCI-186, decreases the capacitance loss fully to baseline. We note in this connection that, among antioxidants, edaravone is one of the most effective to inhibit transition pore openings, thus preventing ischemia reperfusion damage (39) and improving significantly the prognosis for amyotrophic lateral sclerosis patients.

Figures 3D and 3E illustrate the exceptionally powerful OMDE activity of murine BMMs, differentiated for 7 days by hematopoietic growth factor, M-CSF(40). We note that capacitance measurements (not shown) revealed that BMMs develop about 10 times more PM during differentiation. As shown in Fig. 3D, upon PM *opening* by patch clamp with pipettes containing succinate (8 mM), BMMs take in >75% of their PM within 2.5 min. These massive OMDE responses are characterized in Fig. 3E. From left to right, PM loss amounts to only about 10% of the PM when mitochondrial substrates are not included in the pipette. Importantly the cytoplasm contains 10 mM MgATP (and 0.2 mM GTP). Thus, the lack of endocytosis cannot be caused by a lack of nucleotides without mitochondrial substrates.

The OMDE responses are decreased by 65% in BMMs from DHHC5-gene-trapped (GT) mice (3^rd^ bar graph), inhibited >80% by the mitochondria-selective cyclosporine, NIM811 (a blocker of PTPs, (41), fourth bar graph), and inhibited 65% by 2:1 bromopalmitate/albumin complexes (160/80 μM, 5^th^bar graph). The OMDE responses are inhibited similarly when the antioxidant, tiron (42), is included in the pipette at a high concentration (15 mM, 6^th^ bar graph)). This result demonstrates that hydrophilic antioxidants can be effective, albeit only at high concentrations, The responses are inhibited about 70% by when cells are pretreated for 10 min with latrunculin (0.5 μM, 7^th^ bar graph). OMDE responses are effectively absent when β-HB (15 mM) is used as the mitochondrial substrate (8^th^ bar graph), potentially because β-HB has antioxidant activity at or in mitochondrial membranes (43). In complete contrast to β-HB, pyruvate (15 mM) supports OMDE responses that are equally large as succinate responses (9^th^ bar graph). Shown as a red bar graph, endocytic response of BMMs induced by including GTPγS (0.3 mM) in the pipette are fully equivalent to the succinate responses, even in the presence of 10 mM EGTA with no added Ca. Finally, the final bar graph (blue) shows that equivalent OMDE responses are activated by 90 μM Triton-X100 without Ca or substrates, presumably because nonionic detergents at low concentration induce membrane phase separations in complex membranes (12, 44) and thereby promote strongly OMDE.

Results shown in Fig. 4 verify in two further cell types that OMDE responses induced by succinate are dependent on DHHC acyltransferase activity. Using murine lung fibroblasts and acutely isolated peritoneal macrophages from WT mice, OMDE responses in the presence of succinate amounted to 48% and 46% of total PM, respectively. In the former case, responses were reduced to 23% in cells from DHHC5 gene-trapped animals, and in the latter case the endocytic responses were converted to membrane expansion in cells from the gene-trapped animals. In summary, OMDE in macrophages and fibroblasts is very powerful when activated by GTPγS or mitochondrial metabolism of succinate or pyruvate. The endocytic responses are far too large to be accounted for by caveolae (10) and are larger than any descriptions of clathrin/dynamin-dependent endocytosis known to us.

**Figure 4.**
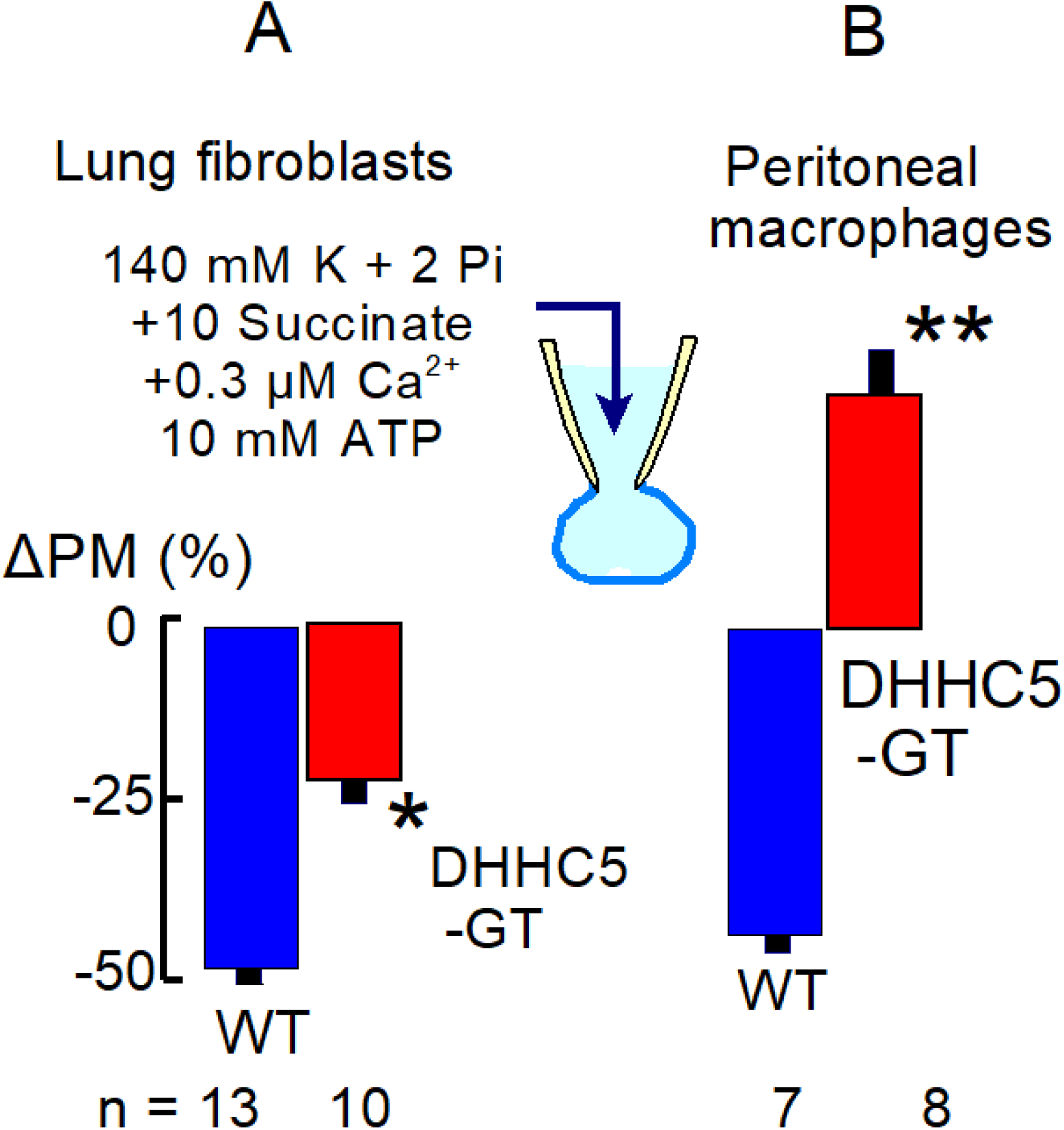
OMDE induced by cytoplasmic succinate in patch-clamped murine lung fibroblasts and acutely isolated peritoneal macrophages under the conditions of Fig. 3E. **Left panel.** OMDE induced by succinate perfusion into lung fibroblasts is reduced by 60% in fibroblasts from animals with gene-trapped DHHC5 (DHHC5-GT). **Right panel.** OMDE induced by succinate perfusion into peritoneal macrophages is blocked and replaced by PM expansion in cells from DHHC5-GT animals.

### OMDE is constitutively active in starved T-REx293 cells but not dependent on DHHC5 or DHHC2

Figure 5 shows initial evidence that OMDE is constitutively active in T-REx293 cells. As described in Methods, cells were incubated in a simple physiological saline solution without serum, substrates, or bicarbonate for 1 h. Thereafter, cells were incubated with 3 kD TexaRed^TM^-lysine-fixable-dextran for 20 min, the PM was labeled with wheat germ agglutinin (WGA) on ice, cells were fixed for 10 min with 4% paraformaldehyde, and nuclei were labeled with Hoechst dye (not shown). Fig. 5A is a typical micrograph of a control cell, showing the overlayed fluorescence of dextran (red) and WGA (green). Fig. 5B shows the time course of dextran uptake, determined from duplicate measurements at the time points indicated after initiating uptake. Dextran uptake saturates within about 10 min, and the apparent time constant of uptake is 7.4 min.

**Figure 5.**
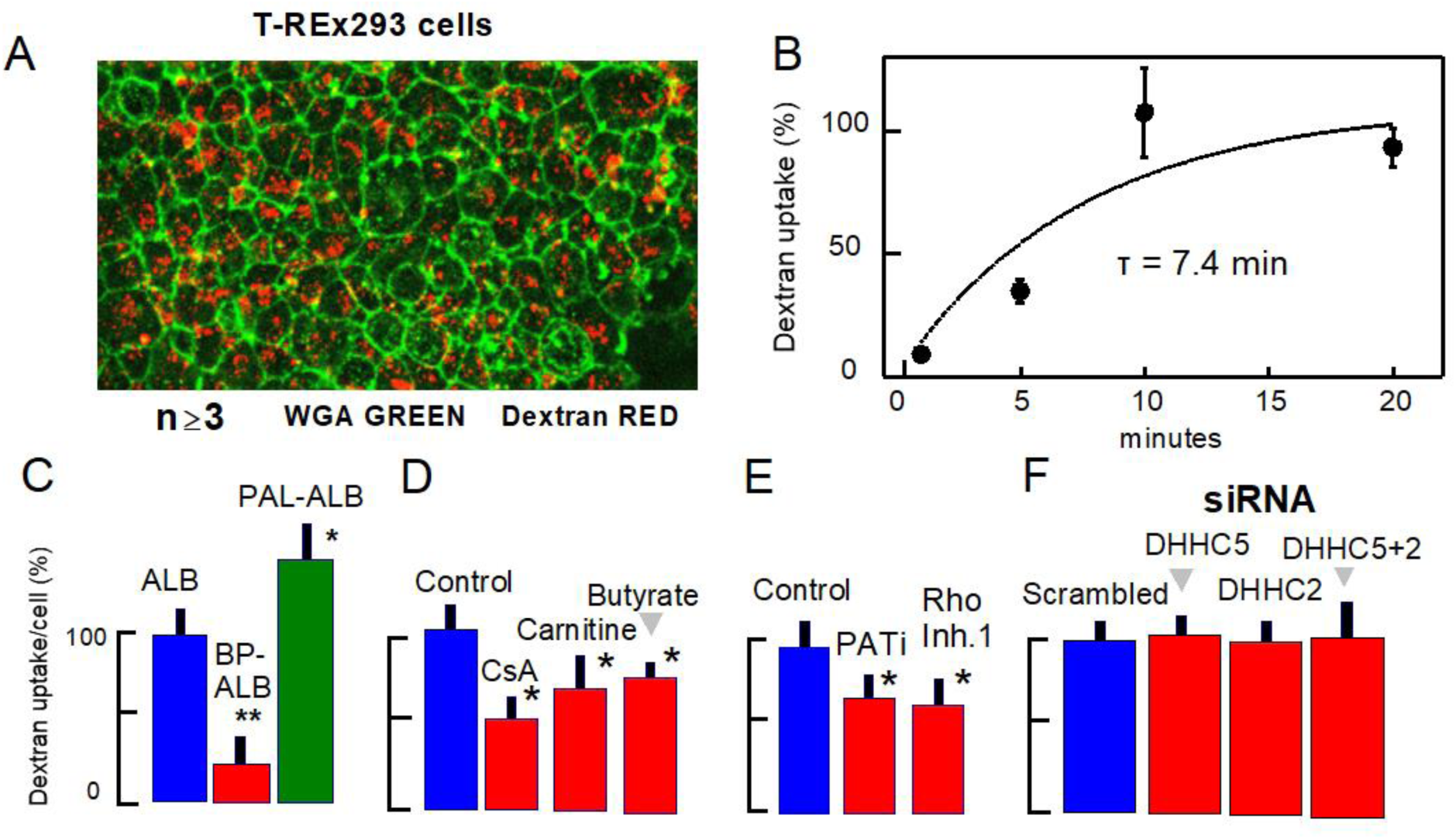
Standard 3kD TexasRed^TM^-dextran uptake assay in a confluent T-REx293 cell culture. **A.** Combined WGA green fluorescence and dextran fluorescence in a culture fixed after 20 min of uptake. Nucleus staining to determine cell numbers is not shown. **B.** The constitutive uptake of dextran saturates over ∼20 min, and the rough time constant is 7.4 min. Therefore, our standard uptake time in subsequent experiments presented was limited to 4 min. **C-F.** Experimental results consistent with a contribution of OMDE under mitochondrial control to constitutive dextran uptake. **C.** Strong inhibition of dextran uptake by bromopalmitate/albumin complexes (0.3/0.1mM, red) versus albumin alone (0.1 mM, blue) and versus uptake in the presence of albumin/palmitate complexes (0.1/0.3 mM, green). **D.** Inhibition of dextran uptake by >50% with cyclosporin A (2 μM), and significant inhibition by carnitine (10 mM) and butyrate (10 mM). **E.** Significant inhibition of uptake by a recently developed DHHC inhibitor (PATi) and by Rho-Inhibitor1. **F.** Failure to inhibit dextran uptake in these substrate- and bicarbonate-free conditions by siRNA knockdown of DHHC5, DHHC2 or combined DHHC5/DHHC2 knockdown.

That OMDE may support constitutive PM turnover in starved T-REx293 cells is supported by the following results (Figs. 5C-5E). Inhibition and stimulation of uptake by bromopalmitate/albumin and palmitate/albumin complexes, respectively, are shown in Fig. 5C. We employed albumin complexes of both compounds to avoid nonspecific membrane effects of these compounds and because albumin effectively delivers these compounds into cells via fatty acid transporters (45). Control uptake (blue) was determined in the presence of 0.1 mM fatty acid-free albumin. Then, in the presence of 3:1 bromopalmitate/albumin complexes (0.3/0.1 mM), which inhibit DHHCs (46), uptake is decreased by ∼70%. In contrast, uptake is increased by 40% in the presence of equivalent palmitate/albumin complexes. As shown in Fig. 5D, dextran uptake is inhibited 60% by 1 μM cyclosporine A (CsA), a nonspecific cyclophilin ligand that is the best documented inhibitor of PTP openings (47), as well as massive endocytosis in patch clamp (37). Subsequent results illustrate inhibition of dextran uptake by metabolites that may shift long-chain acyl-CoA into mitochondria or shift CoA from long-chain acyl-CoA to other CoA metabolites. As shown in Fig. 5D, uptake is decreased significantly in cells incubated for 10 min with carnitine (10 mM) or with butyrate (10 mM). Fig. 5E shows that dextran uptake is decreased by a newly identified DHHC inhibitor (48) (PATi, 10 μM) and is significantly decreased by RhoA inhibition with C3-transferase (Rho Inhibitor I, (49)). We mention in this connection that localized activation of RhoA has recently been shown to activate endocytosis that promotes migration of macrophages in an amoeboidal migration mode (50).

Finally, Fig. 5F presents results of DHHC5 and DHHC2 knockdown experiments using siRNAs, as described in Methods. Verified oligonucleotide sequences were employed to knock down DHHC5 and DHHC2 via transfection with Lipofectamine RNAiMA, and knockdown was verified by Western blotting and qPCR for DHHC5. Knockdown of DHHC2 was verified by qPCR. As apparent in Fig. 5F, neither the knockdown of DHHC5, nor DHHC2, nor knockdown of both DHHCs significantly changed the constitutive uptake of dextran by T-REx293 cells. We conclude therefore that these DHHCs are not essential for constitutive PM turnover in these cells. Other acyltransferases must be involved, if palmitoylation of PM proteins is indeed essential for constitutive endocytosis in the absence of serum, mitochondrial substrates, and bicarbonate.

### Activation of dextran uptake by mitochondrial substrates and bicarbonate

Following up patch clamp studies in Figs. 3 and 4, we next tested if (and when) mitochondrial substrates might significantly affect dextran uptake in T-REx293 cells. To do so, we first included the mitochondrial substrate, acetate (10 mM), in the saline solution employed, and we subsequently included bicarbonate (25 mM, at pH 7.2) to support cell carboxylase activities that might be essential for the effects of mitochondrial substrates. Fig. 6 shows results without bicarbonate. From left to right, Fig. 6A shows that cyclosporin A (1 μM) has a very strong inhibitory effect on dextran uptake in the presence of acetate. Carnitine (10 mM) is mildly inhibitory (Fig. 6B) when compared to patch clamp results. As shown in Fig. 6C, pyruvate (10 mM) inhibits dextran uptake by about 30% in this cellular condition (Fig. 6C), while β-HB (10 mM) inhibits by nearly 80%, similar to cyclosporin A. Inhibition by pyruvate together with cyclosporin A is similar to cyclosporin alone. In Supplementary Fig. S3 we verify that metabolic substrates, specifically pyruvate and β-HB (which generate metabolites activated by CoA), do indeed reduce total long-chain acyl-CoAs by 30 to 50% in HEK293 cells under the conditions of these experiments (i.e without bicarbonate). We show further that carnitine significantly increases long-chain acyl-CoAs, presumably related to a shift of acyl groups into mitochondria.

**Figure 6.**
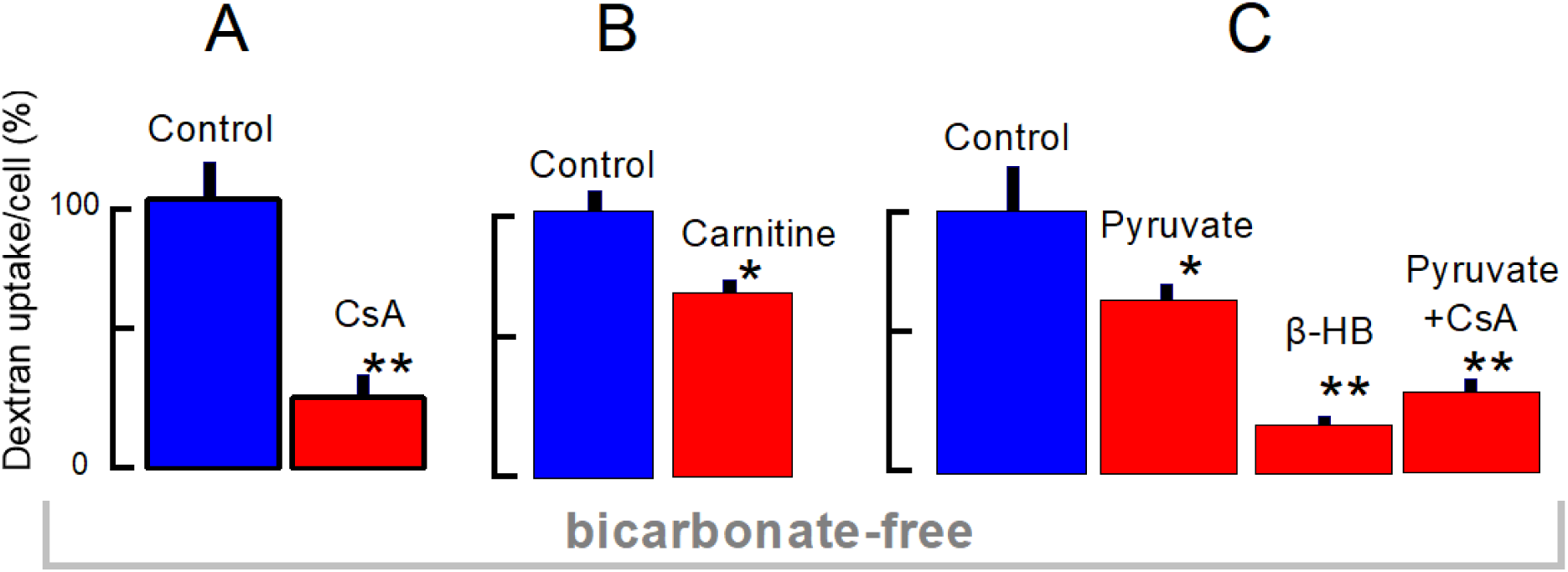
Endocytosis of 3kD TexasRed^TM^-dextran in the absence of bicarbonate and substrates, except for 10 mM acetate. **A**. Dextran endocytosis (4 min) is inhibited 75% by cyclosporin A (CsA, 1 μM). **B.** Dextran endocytosis is inhibited 27% by carnitine (10 mM). **C.** Dextran endocytosis is inhibited 32% by 12 mM pyruvate, 83 % by 12 mM β-hydroxybutyrate, and 73% by cyclosporin A (1 μM) with 12 mM pyruvate.

These results raise an obvious enigma as to why pyruvate is inhibitory for endocytosis in starved, cultured cells but is strongly stimulatory in patch clamp studies. One possible explanation is that pyruvate diffuses rapidly into the cytoplasm during patch clamp experiments, while the cytoplasmic pyruvate concentration likely remains only a small fraction of extracellular pyruvate in experiments with intact cells. Accordingly, the generation of citric acid cycle metabolites from pyruvate may be limited by pyruvate metabolism in intact cells. Of the two pyruvate metabolism pathways, namely decarboxylation to acetyl-CoA and carboxylation to oxaloacetate (51), the lack of bicarbonate in these experiments will limit the pathway to oxaloactetate. As described Fig. 7, the inclusion of bicarbonate (25 mM) in the saline solution employed indeed causes a consistently large stimulation of dextran uptake by pyruvate.

**Figure 7.**
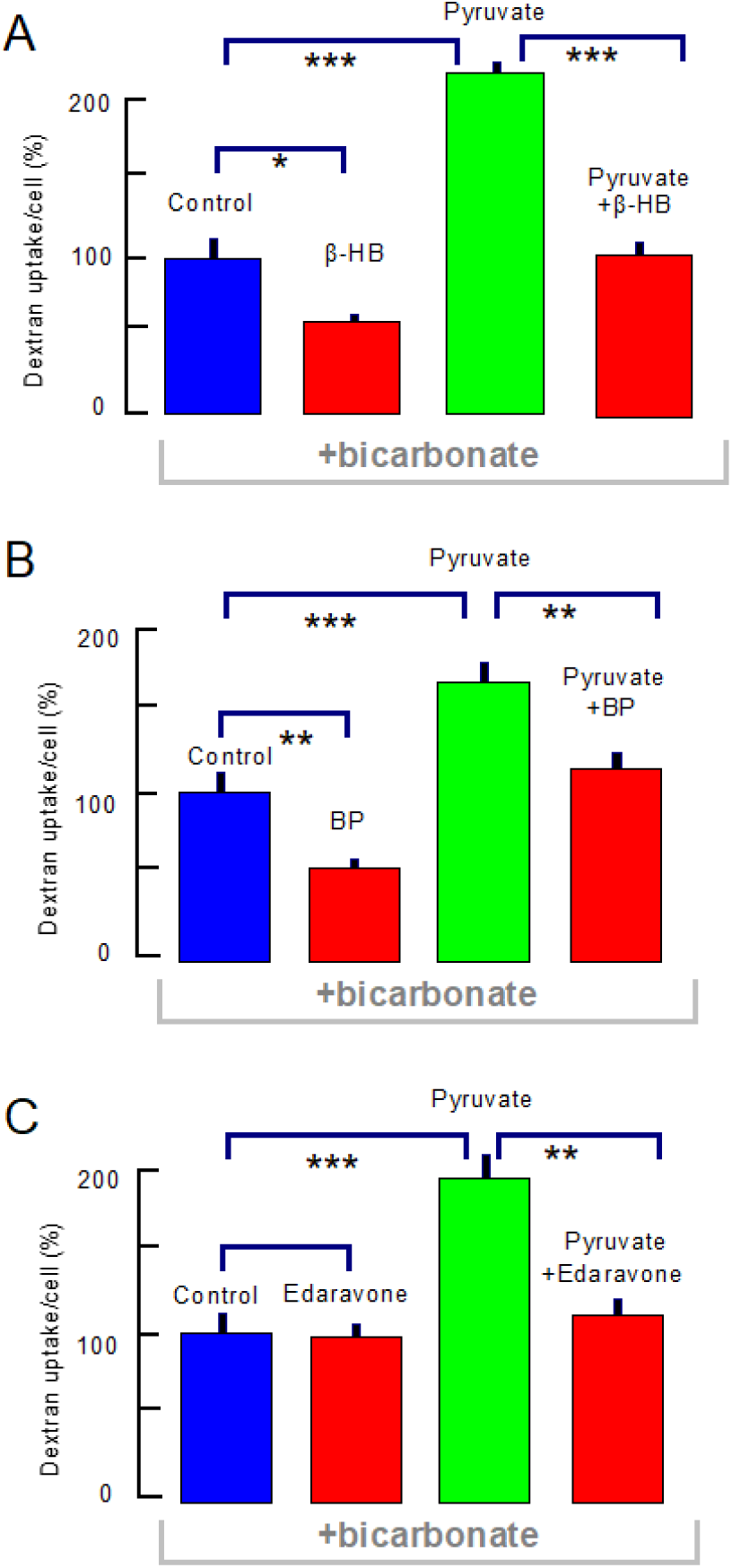
Pyruvate activation of dextran uptake (4 min) in the presence of acetate (10 mM) and bicarbonate (24 mM). **A.** β-HB (12 mM), applied for 30 min prior to the uptake assay, inhibits dextran uptake by 50%, while pyruvate (12 mM) enhances dextran uptake by 2.5-fold. The enhancement by pyruvate is effectively blocked by β-HB (12 mM). **B.** Bromopalmitate/albumin complexes (0.2/0.1 mM), applied for 30 min prior to uptake, decrease constitutive endocytosis by ∼50% and decrease uptake in the presence of pyruvate by 30%. Note that inhibition by β-HB is stronger than the inhibition by bromopalmitate/albumin complexes. C. The antioxidant, edaravone (0.1 mM) is without effect in the control condition but fully blocks the stimulation of dextran uptake by pyruvate (12 mM).

### Strong stimulation of dextran endocytosis by pyruvate and its inhibition by β-HB and bromopalmitate/albumin complexes

In patch clamp studies, pyruvate and β-HB typically have opposite (antagonistic) effects on endocytosis (Fig. 3). Therefore, we compared their effects and interactions in dextran uptake experiments in the presence of acetate and bicarbonate. As shown in Fig. 7A, β-HB (12 mM) reduced dextran uptake by just 50%. In same set of matched cell cultures, pyruvate (12 mM) increased dextran uptake by 2.5 fold. In the presence of β-HB (12 mM), pyruvate (12 mM) increased uptake only to the baseline uptake level. In other words, β-HB effectively blocks the stimulatory effect of pyruvate on endocytosis. As illustrated in Fig. 7B, the inhibitory effects of β-HB are even stronger than those of bromopalmitate/albumin complexes (0.2/0.1 mM) in equivalent experiments. BP complexes inhibit basal dextran uptake similarly to β-HB, about 50%, but pyruvate still increases dextran uptake in the presence of BP to a level that is greater than control uptake. We conclude that β-HB acts with some specificity to block the signaling mechanisms that pyruvate activates to stimulate dextran uptake. As already noted, an antioxidant activity of ketone bodies near or within mitochondrial membranes is one possible explanation (52). As shown in Fig. 7C, we therefore tested the antioxidant, edaravone (MCI-186) at the same concentration (0.1 mM) employed in patch clamp experiments. Different from β-HB, edaravone had no effect on dextran uptake in the control condition. However, edaravone ablated fully the stimulatory effect of pyruvate. This result clearly supports our hypothesis that generation of mitochondrial ROS during pyruvate metabolism is essential for its activation of endocytosis. The result is also consistent with the idea that β-HB acts to inhibit endocytosis by a mechanism besides antioxidant activity, one possibility being a reduction of long-chain acyl-CoAs.

Given that the effects of pyruvate to stimulate OMDE may involve ROS and the opening of PTPs to release CoA, the question is raised whether these mechanisms might cause cells to undergo apoptosis. To test this, we subjected dishes of cells to the protocols that induce endocytosis by pyruvate and then returned the cells to normal culture conditions (i.e with high glucose DMEM medium and 10% serum) for 1 day. After 24 h, we stained cells with trypan blue to test for the presence of dying or dead cells. As illustrated in Supplemental Fig. 4, we found no evidence for damaged or apoptotic cells in cultures that were subjected to the pyruvate protocol.

### Fatty acid-free albumin inhibits dextran uptake at low micromolar concentrations

In our early patch clamp experiments describing massive endocytosis in BHK cells (37), we found that OMDE was inhibited by treating cells with fatty acid free albumin, and we found that OMDE could be restored by restoring fatty acids that were presumably extracted from cells by albumin. Accordingly, we tested to what extent fatty acid free albumin might inhibit constitutive endocytosis in T-REx293 cells. To our surprise, we found that albumin was effective at low micromolar concentrations. Fig. 8A shows results for subjecting cells to 15 μM albumin prior to, but not including, the time of dextran uptake. Fig. 8B shows results for continuous 15 μM albumin exposure, including the dextran incubation period. These protocols are illustrated above the bar graphs in each panel. For results in Fig. 8A, one group of cells was incubated with acetate/bicarbonate-containing saline for 60 min, and then with dextran for 4 min (“1.”). The second group was incubated with saline for 30 min followed by saline with 15 μM albumin for 30 min, and finally with dextran and no albumin for 4 min (“2”). In a third group, albumin (15 μM) loaded with palmitate (30 μM) at a 2:1 ratio was applied for 30 min followed by dextran without albumin (“3”). As summarized by bar graphs, treatment with albumin reduced dextran uptake by 57%, whereas treatment with palmitate-loaded albumin reduced uptake by only 38%. Fig. 8B shows the equivalent results when albumin was administered for 10 min followed by dextran for 10 min with albumin or palmitate-loaded albumin. In this protocol, dextran uptake was reduced 40% by albumin, and the presence of palmitate entirely precluded inhibition by albumin. As described next, it appears that FA-free albumin indeed can in fact decrease palmitoylations of PM proteins that may be involved in endocytosis, an effect that would be consistent with localized FA and acyl-CoA turnover at the cytoplasmic face of the PM.

**Figure 8.**
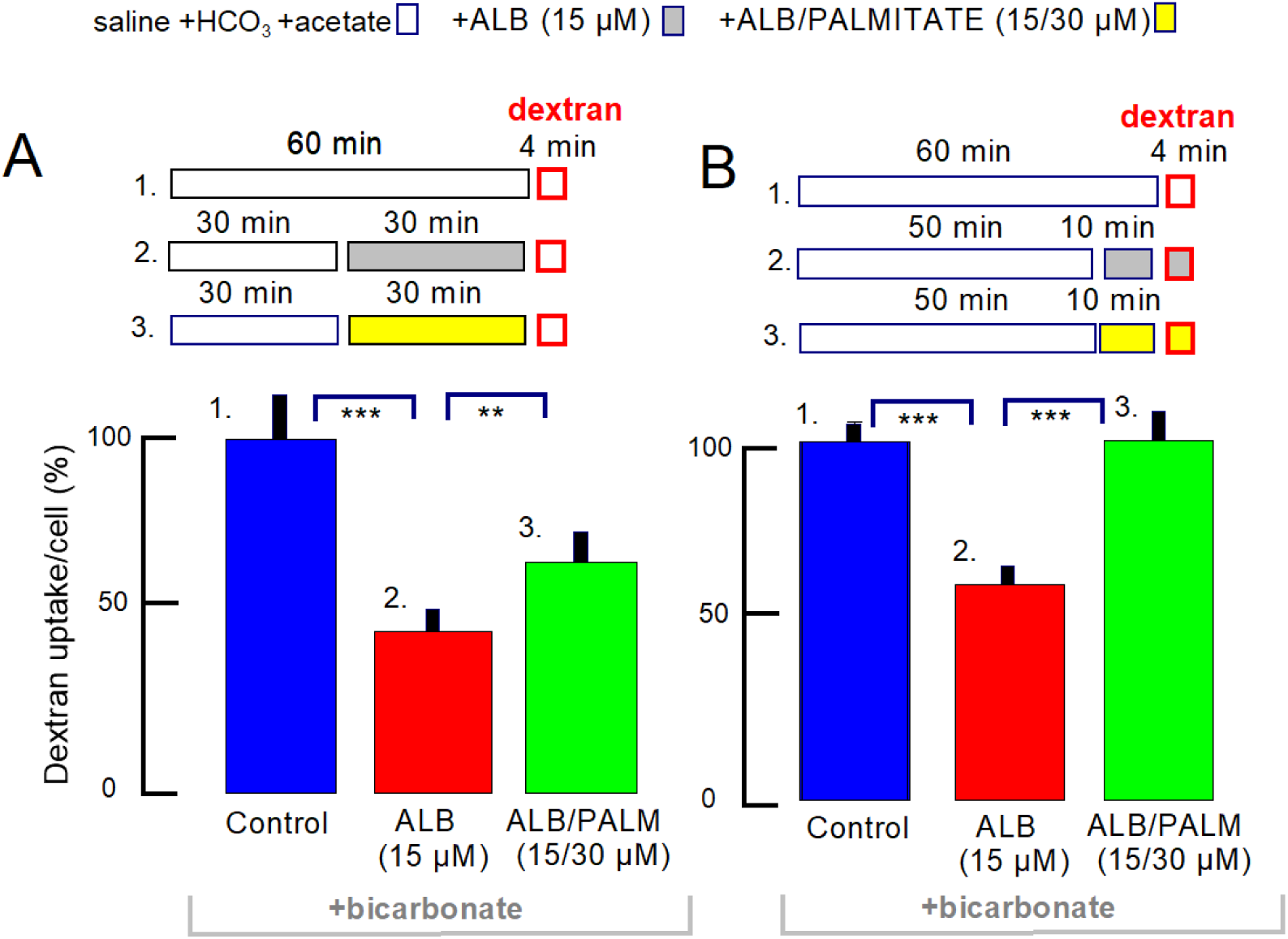
Dextran uptake (4 min with acetate and bicarbonate) is differentially inhibited by low (15 μM) concentrations of FA-free albumin. **A.** As indicated above the bar graphs, control cells were incubated for 60 min with acetate/bicarbonate-containing saline followed by 4 min dextran uptake (“1”). The second (“2”) and third (“3”) sets of cells were incubated with saline for 30 min followed by 30 min with 15 μM albumin or 15 μM albumin with 30 μM palmitate. Dextran uptake was then initiated for 4 min in the absence of albumin and palmitate. Albumin exposure inhibited subsequent dextran uptake by 54%. **B.** As indicated above the bar graphs, control cells were incubated in bicarbonate/acetate-containing saline for 60 min, followed by dextran uptake for 4 min (“1”). Additional sets of cells were incubated for 50 min with saline followed by 10 min with albumin (15 μM) or albumin with palmitate (15 μM/30 μM). Dextran uptake was then initiated for 4 min with albumin or albumin with palmitate, as indicated. The inhibitory effect of albumin alone amounted to 39%, and the presence of palmitate with albumin ablated the inhibitory effect of albumin.

### Inhibition of PM palmitoylations by carnitine, β-HB and FA-free albumin

Next, we tested whether key protocols that inhibit or stimulate endocytosis in fact modify palmitoylations of membrane-associated proteins. To do so we incubated 10 cm dishes of T-REx293 cells equivalently to the protocols employed for uptake assays. Cells were incubated with the saline solution containing acetate and bicarbonate for 1 h. Thereafter, cells were incubated with either 10 mM carnitine + 10 mM β-HB or with 15 μM FA-free albumin for 20 min. Separate groups of experiments were performed with addition of 10 mM pyruvate, harvesting three dishes each after 2 and 10 min. Cells were then processed as described in Methods. Free cysteines were blocked immediately upon cell harvesting, a crude membrane fraction was isolated, proteins were deacylated with hydroxylamine, and proteins with free cysteines were pulled down thiol reactive agarose beads (G-Biosciences.com)(53).

Figure 9A shows the average changes of protein counts for the four treatments examined with control counts normalized to 100%. As described in Methods, we normalized results for each protein sample to the counts obtained for caveolin-1, whose prevalence never changed significantly in the deacylated protein samples. Results shown were limited to the top 1200 protein species identified in the deacylated samples, more than 90% of which were enriched more than two-fold in hydoxylamine-versus TRIS-treated samples. As shown in the bar graphs, 20 min exposures to carnitine and β-HB decreased protein counts on average by 52% (p<0.001), which 20 min exposure to FA-free albumin (15 μM) decreased average protein counts by 50% (p<0.001). In contrast, treatment of cells with pyruvate (10 mM) for 2 min increased average deacylated protein counts by 69% (p<0.001) and the continued exposure to pyruvate (10mM) for 8 further minutes increased the average deacylated protein counts to 209% of control counts.

**Figure 9.**
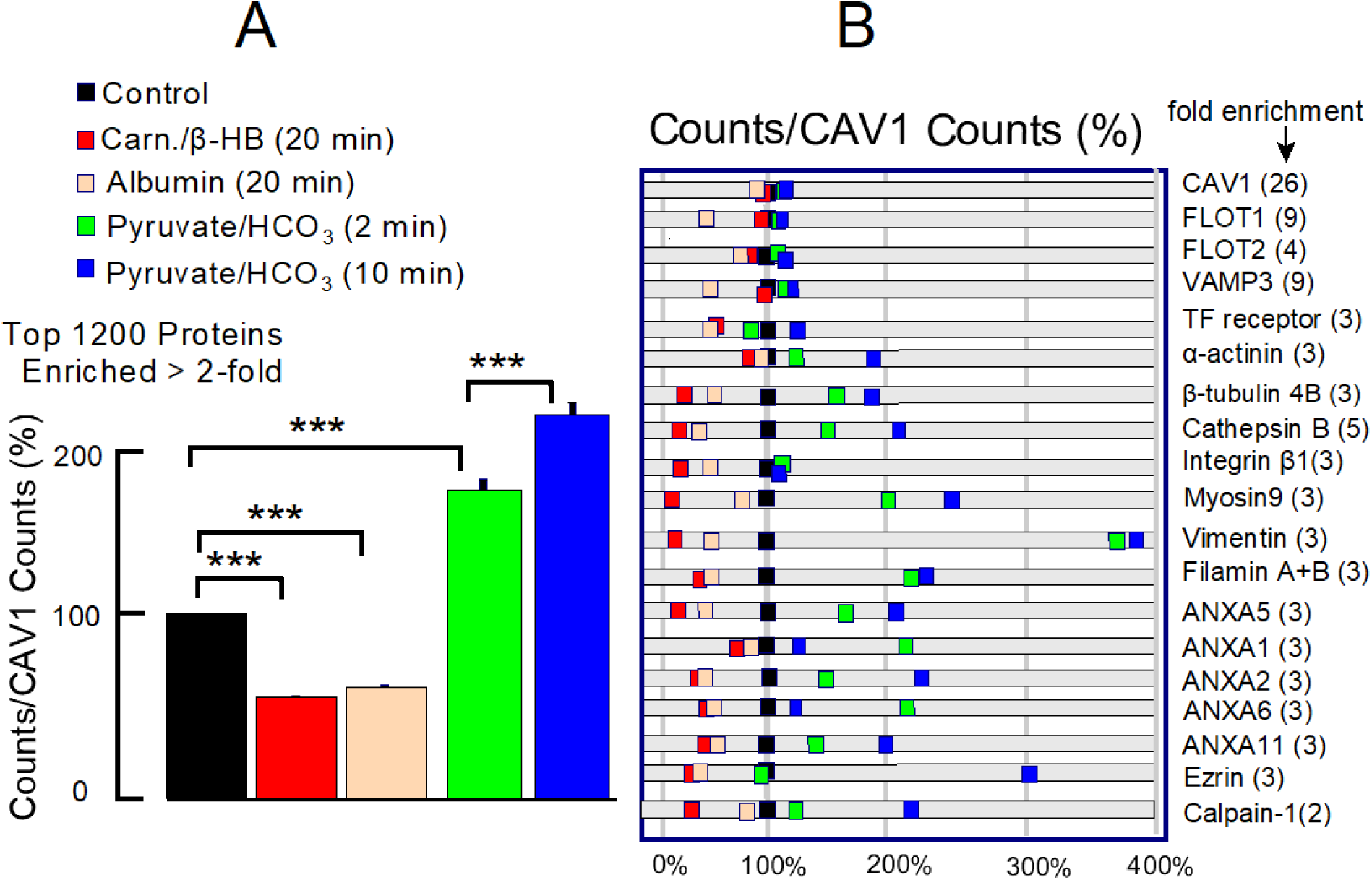
Biochemical and cell biological evidence for selective role of palmitoylation in OMDE. As indicated by red and tan rectangles, cells were incubated in substrate- and bicarbonate-free solution for 1 h, followed by 20 min incubation with either carnitine (10 mM) + β-HB (10 mM) or with FA-free albumin (15 μM). As indicated by green and blue rectangles, cells were incubated with acetate (10 mM) and bicarbonate (24 mM) for 1 h, followed by incubation with pyruvate (10 mM) for 2 or 10 min respectively. Results from each sample were normalized to the protein counts for caveolin-1, which were not significantly changed by the treatments. Results for all experiments were normalized to the respective control results. **A.** Counts for the top 1200 proteins in hydroxylamine-treated samples. Carnitine/β-HB treatment reduces average counts by 47%, albumin treatment reduces counts by 39%, and pyruvate in the presence fo acetate and bicarbonate increases counts by 69 and 100% at 2 and 20 min, respectively. **B**. Results for selected proteins that are represented in the SwissPalm data base multiple times and verified by three methods. Many proteins that did not change are known to be palmitoylated at multiple sites (e.g. flotillin-1, flotillin-2,and VAMP3.

Selected results from the proteomic analysis are shown in Fig. 9B. In brief, the palmitoylation of many PM proteins, confirmed multiple times in the SwissPalm palmitoylation database with at least 3 different methods (https://swisspalm.org), are decreased. Notably, many of the proteins that follow the average pattern described in Fig. 9A, while identified by multiple methods in the SwissPalm data base, do not have high stringency sequences for palmitoylation as represented in the CSS-Palm 4.0 palmitoylation predictor program (see (54)). From top to bottom, caveolin-1 prevalence did not change in this experiments, presumably indicating that at least on palmitoylation remains with all treatments employed. Flotillin-1 palmitoylation decreased significantly with albumin, but not other flotillin result was striking. Examples of proteins that mirrored the average results, or showed still larger changes, included the transferrin receptor, β-tubulin 4B, cathepsin B, integrin β1, myosin-9, vimentin, filamins A and B, annexins-5, −1,-2,-6 and −11, and calpain-1. The identification of multiple Annexin proteins in this assay may be relevant to OMDE activation by Ca (31), and the cytoskeleton-associated proteins identified, including vimentin and filamins, may be relevant to the actin-dependence of OMDE.

### Reciprocal effects of palmitoylation on uptake of cholera toxin B (CTB) and transferrin (TF)

Knowing that multiple biological interventions that inhibit or stimulate OMDE also would appear to inhibit or stimulate palmitoylations similarly, we next examined the effects of one of those treatments (carnitine+β-HB) on endocytosis defined by two classical membrane markers, CTB and TF. While neither of these probes is entirely specific, CTB binds selectively GM1 gangliosides and is internalized primarily by clathrin/dynamin-independent mechanisms. We note however that CTB is also taken up to some extent by clathrin endocytosis (55, 56) in addition to ‘raft’ endocytosis. TF, on the other hand, is internalized primarily by classical clathrin-dependent endocytosis (57). Using a 2 min uptake assay under substrate- and serum-free conditions to define constitutive uptake, the uptake magnitudes were similar for the two probes (see micrographs in Figs. 10A and B, and composite results in Figs. 10C and 10D). Nevertheless, uptake of CTB was decreased ∼40% by the carnitine/β-HB treatment (p<0.01) while TF uptake was significantly increased (p<0.01). Regarding the increase of TF uptake, it was described already in 1990 that transferrin palmitoylation hinders clathrin endocytosis and that depalmitoylation promotes iron uptake (58). Overall, these results verify that the major two pathways of endocytosis, as defined here, are affected differently by interventions that affect formation of ordered membrane domains. The results are consistent with OMDE mediating much of the CTB uptake occurring in T-REx293 cells. While OMDE contributes to basal PM turnover in T-REx293 cells, a substantial contribution of clathrin endocytosis is also indicated.

**Figure 10.**
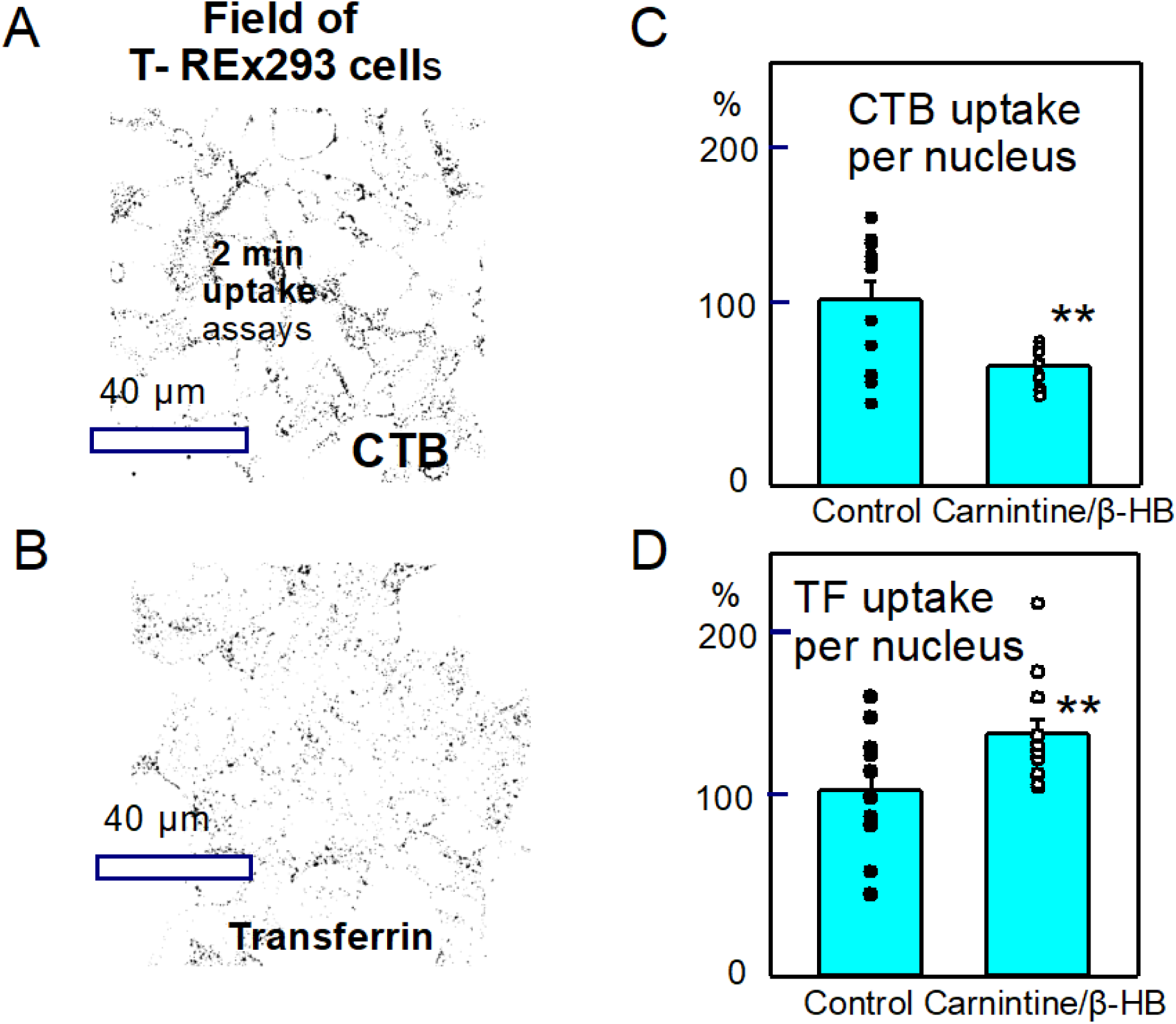
Effects of carnitine (10 mM) + β-HB (10 mM) on the uptake of cholera toxin B (CTB) and transferrin for 2 min in T-REx293 cells. Aurox near-confocal imaging. **A,B.** Negative images of the micrographs of CTB and transferrin uptake. **D,E.** Effects of treating cells with carnitine+β-HB for 10 min. TB uptake is decreased is by 37% (p<0.02) while TF uptake was increased by 22% (p<0.01).

### STED microscopy reveals dimensions of endocytic vesicles formed by constitutive OMDE

Finally, to address the membrane structures formed during OMDE, we initiated super-resolution optical studies with STED microscopy (59). Using 10 kD CF640R-dextran to monitor fluid phase endocytosis, Fig. 11A shows a representative micrograph for 2 min dextran uptake in T-REx293 cells. Within the 2 min, 514 puncta form with diameters described in histograms in Figs. 11B and 11C. Converting the surface areas of particles to spherical areas, the uptake constitutes a loss of ∼110 μm^2^ of PM. This corresponds to ∼1 pF or ∼1% of the PM of a large T-REx293 cell. The histograms of puncta diameters appear to have two components in control cells (Fig. 11B). To acutely inhibit OMDE, we treated cells simultaneously with carnitine (5 mM) and β-HB (5 mM) for 30 min. The histogram from treated cells, shown in Fig. 11C, is paired experimentally with the control histogram in Fig.11 B. The treatment employed results in a decrease of average vesicle diameter from 114 to 95 μm. and a decrease of mean vesicle diameter from 94 to 71 nm (p<0.001). Cell morphology was stable, and the number of particles in the size range from 85 to 130 nm was significantly decreased. Assuming that OMDE is primarily blocked in these experiments, rather than clathrin endocytosis, we conclude that the vesicles formed by OMDE are on average larger than vesicles formed by clathrin endocytosis but substantially smaller than vesicles formed by macropinocytosis (60).

**Figure 11.**
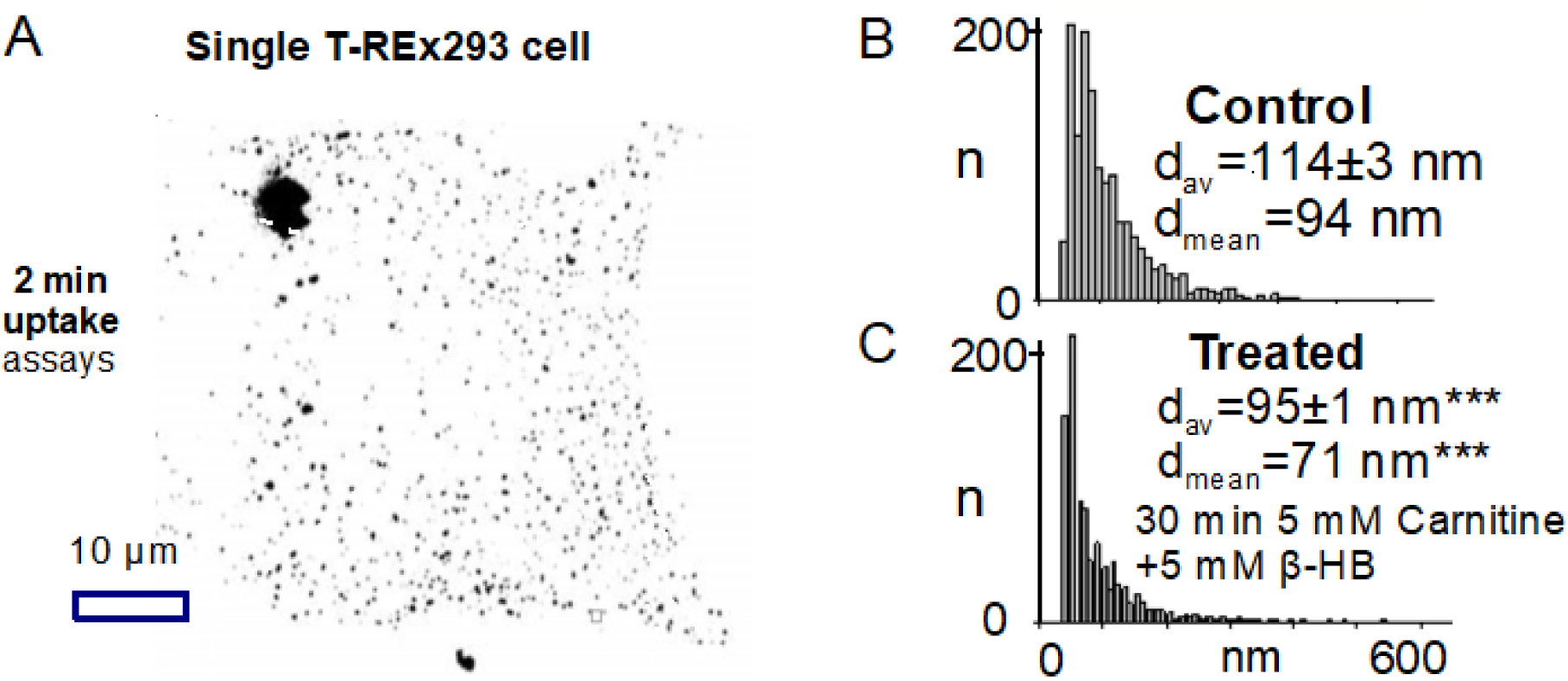
Vesicle dimensions determined via STED microscopy of CF640-dextran uptake for 2 min. **A.** Negative image of a single T-REx293 cell, visualized with STED microscopy after CF640R dextran incubation. Cells were serum, substrate and bicarbonate-free for 20 min. STED resolution was determined to be ∼30 nm, thereby allowing **a**ccurate quantification of endocytic vesicle dimeters. **B,C.** Histograms of puncta diameters under control conditions and after treatment for 30 min with carnitine (5 mM) +/β-HB (5 mM). The histograms reveal a highly significant decrease of average and mean vesicle sizes. Particle counts within the diameter range of 80 to 140 nm are significantly decreased (p<0.01). The loss of puntae amounts to ∼50%.

### Mitochondria-dependent OMDE does not contribute to constitutive endocytosis in BMMs

Finally, Figure 12 presents our analysis of constitutive endocytosis in BBMs, i.e. in the absence of serum, substrates, and bicarbonate, indicating that OMDE is inactive in the basal state of these cells. Fig. 12A is an overlay from a representative assay of nucleus outlines and particle outlines in an Aurox field of view. There are 120 nuclei and on average 30 particles per cell, about 3-fold fewer particles per cell than found in T-REx293 cells under these conditions and using the same ImageJ analysis parameters. Fig. 12B shows that bromopalmitate/albumin complexes (0.3/0.1 mM) have no effect on dextran uptake in experiments carried out identically to those for T-REx293 cells in Fig. 5. Palmitate/albumin complexes (0.3/01 mM) increase uptake at most marginally. Fig. 12C shows that a high cyclosporin A concentration (2 μM) has no effect on uptake, and Fig. 12D illustrates the insensitivity of uptake to addition of β-HB (10 mM) and pyruvate (10 mM). Finally, shown in Fig. 12E, dextran uptake was not different under these conditions in BMM cultures from mice lacking DHHC5. From these results, we conclude that the mechanisms mediating massive endocytosis in BMMs, as observed during patch clamp with mitochondrial substrates or GTPγS (Fig. 3B), are effectively inactive in the absence of serum, substrates, and bicarbonate.

**Figure 12.**
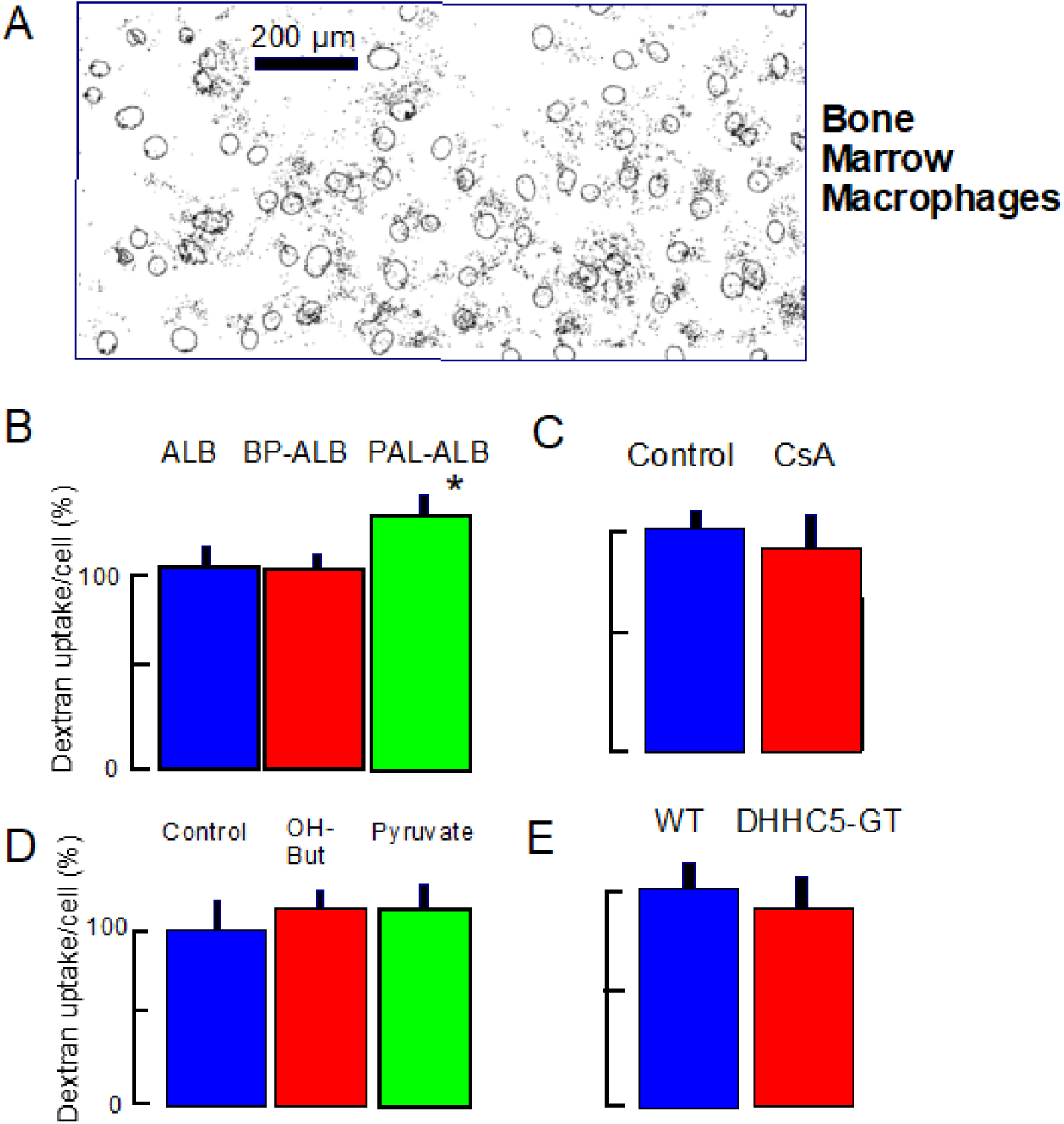
Uptake of TexasRed^TM^-dextran in BMMs, differentiated in cell culture for 7 days. Uptake was performed in the absence of serum, metabolic substrates, and bicarbonate. Particle counts per cell ranged from 30 to 50% of those obtained in T-REx293 cells. **A.** Typical outlines of nuclei (Hoechst dye) and endocytic particles (puncta) determined with Aurox near-confocal imaging after 20 min of TexasRed^TM^-dextran uptake. **B.** In contrast to T-REx293 cells under the same condition, bromopalmitate/albumin complexes are not inhibitory, and stimulatory effect of equivalent palmitate/ albumin complexes are very small. **C.** In contrast to T-REx293 cells under the same condition, cyclosporin A (CsA) is without effect on dextran uptake. **D.** In contrast to T-REx203 cells, β-HB (10 mM) and pyruvate (10 mM) do not inhibit constitutive dextran uptake. **E.** Dextran uptake under these conditions is not less in DHHC5 gene-trapped BMMs than in WT BMMs.

## Discussion

In this article we verify that three forms of “activated” OMDE are supported by and/or require the acyltransferases, DHHC5 and DHHC2. We have compared constitutive endocytosis in T-REx293 cells and murine BMMs, and the outcomes are very different. Without serum, substrates, or bicarbonate, T-REx293 cells have a large endocytic flux that is consistent with OMDE under mitochondrial control (Fig. 5), and equivalent experiments in BMMs are entirely negative (Fig. 12). In contrast to “activated” OMDE, we find no involvement of DHHC5 or DHHC2 acyltransferases in constitutive OMDE in T-REx293 cells (Fig. 5F). We have shown that mitochondrial metabolites, namely succinate, pyruvate and β-HA, can markedly modify OMDE, and that classical clathrin endocytosis can be affected in a reciprocal manner (Fig. 10). Remarkably low FA-free albumin concentrations can inhibit constitutive OMDE in parallel with an apparent decrease of PM protein palmitoylations (Figs. 8 and 9), while pyruvate stimulation of OMDE appears to be accompanied by increased palmitoylations of many PM-associated proteins (Fig.9). Finally, we have analyzed the physical basis of constitutive OMDE via STED microscopy in T-REx293 cells (Fig. 11). Endocytic vesicles formed have diameters of 90 to roughly 140 microns, They are thus larger than average vesicles formed via coated pits (i.e. clathrin endocytosis) but substantially smaller than vesicles formed by macropinocytosis (60). We discuss the Results roughly in the order of their presentation.

### OMDE activated by G-proteins and oxidative metabolism in T-REx293 cells and BBMs

We have documented in Figs. 3 and 4 the extraordinary power of OMDE when activated by cytoplasmic GTPγS or mitochondrial substrates in T-REx293 cells and murine BBMs. The occurrence of massive endocytosis in the presence of a high (0.2 mM) concentration of GTYγS, which cannot be hydrolyzed by dynamins, makes the involvement of dynamin GTPase very unlikely (31). While we have not defined which G-proteins are essential, the involvement of multiple small G-proteins in clathrin/dynamin-independent endocytosis has been well established for decades (61). Inhibition of the small G-protein RhoA is clearly inhibitory (Fig. 5E). We demonstrate in Fig. 2 that OMDE activated in both T-Rex293 cells and BMMs is dependent on DHHC2 and/or DHHC5, as well as actin cytoskeleton. In the case of T-REx293 cells we have used siRNA of these two DHHC enzymes, known to reach the PM (34, 35), to demonstrate a dependence of GTPγS-activated OMDE on acyltransferase activity. In the case of BMMs, we have employed cells from DHHC5 gene-trapped animals (20) to demonstrate dependence of succinate-activated OMDE on acyltransferase activity. For lung fibroblasts and acutely isolated peritoneal macrophages, we have also demonstrated a dependence of succinate-activated OMDE on DHHC5 with cells from gene-trapped animals (Fig. 3A).

Concerning actin-dependence, endocytosis induced both by GTPγS and by succinate metabolism is decreased more than 60% by latrunculin (Figs. 3B and 3E). This contrasts to our previous studies of massive endocytosis induced by Ca elevations in which latrunculin was without effect (13, 31). One obvious difference between the different forms of OMDE is that Ca elevation initially causes a large PM expansion in most cell types, which may relax PM tension (31). Consistent with this interpretation, the block of OMDE in T-REx293 cells by latrunculin can be fully overcome by cell shrinkage, employing either hypotonic cytoplasmic solutions or negative pressure applied to the pipette tip (Fig. 3B). Both of these interventions will with little doubt decrease lateral membrane tension. It will now be of considerable interest to test whether inhibitory effects of actin disruption on ‘bulk endocytosis’ in neurons (62) are also dependent on lateral PM tension.

We have suggested previously that ROS generation by mitochondria is a major signaling factor that promotes massive endocytosis via OMDE (37). ROS may act through many signaling mechanisms, including the activation of small G-proteins (63) such as RhoA (64). In addition, however, our work suggests that the opening of mitochondrial PTPs (permeability transition pores) is critical, namely to release mitochondrial metabolites, in particular CoA, to the cytoplasm (37). These suggestions are supported presently by the effective block of succinate-activated OMDE in BMMs by the mitochondria-specific cyclosporin, NIM811, and the inhibition observed with two antioxidants, edaravone and tiron (Fig. 3). Admittedly, tiron is not specific, as it chelates iron and other multivalent metals (42). We point out in this connection that attempts to block PTP openings with antioxidants have often had mixed or complex outcomes (e.g. (65, 66)). The effectivity of edaravone in both patch clamp and dextran uptake assays (Figs. 3 and 7C) is impressive, and this is consistent with the clinical application of edaravone in ALS (67).

The consistently small magnitude of OMDE that occurs with β-HB, and the ability of β-HB to suppress pyruvate activation of OMDE (Fig. 7A), may be relevant to its beneficial effects in several degenerative disease models (e.g. (68)). The fact that cytoplasmic pyruvate, like succinate, strongly activates OMDE in both T-Rex293 cells and in BBMs was surprising, since within the cytoplasm pyruvate likely has antioxidant functions (e.g.(69)). Presumably, ROS generation at or within the inner mitochondrial membrane is critical for PTP openings and the progression of OMDE. In the simplest case, β-HB may function as a membrane-near antioxidant at the critical mitochondrial sites, presumably the immediate vicinity of cytochrome oxidases. Importantly, we have shown that β-HB can reduce long-chain acyl-CoAs in cells (Fig. S3), an effect that rationally explains why β-HB is more effective than edaravone in reducing OMDE in the absence of pyruvate (FIg. 7).

The speed and magnitude of OMDE activated by succinate and pyruvate in BBMs (Fig. 3) are unprecedented for any form of biological endocytosis known to us. On average 77% of the PM is removed within 2 to 3 minutes. Since these responses can occur even when cytoplasmic Ca is heavily buffered (10 mM EGTA with 3 mM Ca (Fig. 3B)), Ca elevations are definitively not required. As already stressed, the generation of ROS and PTP openings are both implicated for succinate-induced OMDE. The fact that endocytosis induce by pipette perfusion of GTPγS is very similar in magnitude (and speed) indicates that G-proteins besides dynamins are very likely involved.

Nonionic detergents are long known to cause phase separations of complex membranes at low concentrations (see (44)). Thus, the induction of very large endocytic responses low concentrations of TritonX-100 in BMMs is further consistent with our hypothesis that growth of ordered domains drives the endocytic responses to mitochondrial substrates and GTPγS. These results support further the notion that phase separations of plasma membranes involves the lateral rearrangement of both lipids and PM proteins. By involving PM proteins, especially palmitoylated proteins with large cytoplasmic domains, OMDE can likely internalize larger fractions of the PM and generate smaller vesicles than would be expected for lipid rafts, per se. (70).

### The constitutive roles and regulation of PTPs

While it is firmly established that mitochondrial PTP openings are associated with mitochondrial depolarization, and eventually the initiation of cell death programs (71), the physical identity of PTPs and their signaling roles in normal cell physiology remain controversial (72, 73). That PTPs can open transiently in a so-called ‘flickering’ mode during normal cell physiology is well established, and such openings are in general thought to have a low conductance that is adequate for depolarization, release of calcium, and generation of ‘flashes’ of mitochondrial ROS. Our hypothesis, however, that PTPs can release significant amounts of CoA physiologically requires that physiological PTP openings generate a significant permeability to relatively large solutes. In fact, studies employing calcein release to monitor PTP openings do indeed document significant solute release in a spontaneous and maintained fashion from subpopulations of mitochondria (74, 75), and such openings have been implicated to initiate signaling that reprograms the development of certain cell types (76). In a larger biological perspective, solute release associated with electrical spiking is documented to occur in bacteria (77). It is as least plausible that nonspecific solute release represents an evolutionarily primitive form of intercellular signaling (e.g. (78)).

A strong limitation of our work on OMDE up to now has been the fact that most experiments involved patch clamp, an experimental model that does not readily allow biochemical studies. The protocols described in this study, which activate and modify OMDE in cells growing in dishes, open a wide range of new experimental possibilities to study OMDE. The strong effects of cyclosporin A and pyruvate on dextran uptake in cells cultured with bicarbonate (Figs. 6 and 7), for example, provide new ways to test whether the generation of long-chain acyl-CoAs and cysteine palmitoylations really underlie the initiation of OMDE.

### Constitutive activity of OMDE is highly variable between cell types

Often, attempts to define specific endocytic mechanisms and their involvement in PM turnover employ molecular biological means to overexpress or to delete individual endocytic proteins. The caveat then is that cells may adapt and compensate genetic interventions in the time frame required to induce protein changes. We elected therefore, for the purposes of this article, to examine acute interventions that with reasonable confidence can indicate which mechanisms are active. To test whether OMDE under mitochondrial control is constitutively active in T-REx293 cells, we initially cultured cells without serum, substrates, or bicarbonate for 1 h in a physiological saline solution. In this environment (Fig. 3), we found that PM turnover was strongly decreased by bromopalmitate/albumin complexes and could be increased by albumin/palmitate complexes. These results are not highly specific, but together with other results do suggest that palmitoylations of PM proteins support basal PM turnover. Turnover is decreased by >50% by a cyclosporin (cyclosporin A) that suppresses PTP openings. Of course, cyclosporin A also inhibits the phosphatase, calcineurin (79), although rather high concentrations appear to be required to definitively decrease clathrin/dynamin-dependent endocytosis (80).

We have shown previously (37), as well as in the present study (Fig. 3E), that the mitochondria-specific cyclosporin, NIM811 (81), effectively inhibits OMDE. Furthermore, constitutive PM turnover is decreased by a recently developed DHHC inhibitor, PATi (48). Basal PM turnover is significantly decreased by carnitine, expected to shift acyl-CoA from the cytoplasm into mitochondria via the intermediate, acyl-carnitine. In addition, basal PM turnover is inhibited significantly by inhibiting the small G-protein, RhoA. consistent with evidence that multiple forms of OMDE are dependent on small G protein activity. In the presence acetate (10 mM) but no bicarbonate, inhibition of dextran endocytosis by cyclosporin A is particularly strong, as is the inhibition by hydroxybutyrate. Inhibition by carnitine is significant, and pyruvate itself is just as inhibitory (Fig. 6). Importantly, we have verified that both β-HV and pyruvate indeed decrease long-chain acyl CoAs under these specific conditions. The results together clearly support the idea that OMDE under mitochondrial control contributes to constitutive PM turnover in T-REx293 cells. Impressively, all of these same experiments yield negative results for constitutive PM turnover in differentiated BMMs, even though OMDE, when activated, can be much more powerful in BMMs than in T-REx293 cells.

As noted in the Introduction, the present study was initiated in response to an article suggesting that constitutive PM turnover in HeLa cells occurs exclusively via conventional clathrin/dynamin-dependent endocytosis (9). We were surprised at this result, given that OMDE can be extraordinarily powerful when activated. But the clear outcome of the present study is that OMDE may or may not play significant roles in basal PM turnover. In strong contrast to results for T-REx293 cells, experiments using BMMs do not suggest any role of OMDE in basal PM turnover (Fig. 12), defined as here as PM turnover that occurs without serum, substrates, or bicarbonate. Clearly, the role of OMDE in constitutive PM turnover is highly variable and certainly will depend on signaling status of cells in culture. In the present study it is also notable that constitutive endocytosis in T-REx293 cells dependent on fatty acids and several factors that are shown to change palmitoylations, but it is not dependent on the presence of the DHHC2 or DHHC5 acyltransferases (Fig. 5F). Given that cells contain more than 20 DHHC isoforms (82), it would not be surprising if different DHHC enzymes are critical in different cell conditions. Assuming that coenzyme A and long-chain acyl-CoAs occur at relatively low concentrations in the conditions of these measurements, DHHC enzymes with relatively high affinities for substrates might likely be required.

### New support for the existence of palmitoylation-dependent endocytosis

Our experiments addressing palmitoylations on a proteome-wide scale (Fig. 9) now lend new support to the hypothesis that palmitoylations of PM proteins importantly promote OMDE (14). In fact, the adhesion receptor, CD44, and the scavenger receptor, CD36, were both shown 20 years ago to internalize via ordered membrane domains (rafts) in a palmitoylation-dependent manner (83, 84). By demonstrating apparently wide-spread changes of palmitoylations, which correlate qualitatively substrate-dependent changes of OMDE (Fig. 9), the present work supports the idea that palmitoylation-dependent OMDE is likely to be a physiologically important mechanisms of PM turnover. In some cases, interventions that are expected to block the OMDE pathway outlined in Fig. 1 constitute more that 50% of PM turnover. In general, inhibitory interventions reduce constitutive endocytosis in T-REx293 cells by about 50% (Figs. 5-8 and Fig. 10), and it is very possible that larger effects of cyclosporin A and bromopalmitate may be nonspecific. Importantly, and perhaps most definitive, our analysis of dextran uptake via STED microscopy (Fig. 11) is consistent with palmitoylation-dependent OMDE and conventional endocytosis making up approximately equal shares of constitutive PM turnover in this cell type. Of course, this does not change the fact that OMDE can become much more powerful when activated.

As noted in the Introduction and documented in Results, OMDE involves the ordering of both lipids and membrane proteins into domains and can also involve the actin cytoskeleton.Thus, OMDE does not rely only on membrane ordering to drive vesiculation. It seems likely at this time that the presence of different protein cargos initiates OMDE with different physical characteristic. In the form of OMDE dubbed ‘CLIC’, for example, the palmitoylated ‘raft’ protein, CD44, is a major cargo, and endocytic vesicles take on a tubular form that may be determined both by CD44 and the branching of actin cytoskeleton that appears to contribute to the endocytic process (16, 85). Up to now, visualizing endocytic structures with STED microscopy, we have not clearly observed tubular structures. Accordingly, it will be important to test whether the overexpression of different cargo proteins can dictate how OMDE develops and which structural features can be influenced by the expression of different cargos.

The present body of work provides a new basis to analyze OMDE using coordinated patch clamp and cell culture/biochemical approaches. An especially surprising result, which requires further experimental attention, is that low concentrations of FA-free albumin substantially decrease OMDE in parallel with a decrease PM protein palmitoylations (Fig. 8). Two interpretations of these results are still possible. On the one hand, albumin is known to directly interact with membranes, in particular cholesterol-rich membranes (e.g. (86)). It remains possible therefore that FA-free albumin might induce changes of membrane structure, possibly PM ordering, that either inhibit the palmitoylation of PM proteins or promote the depalmitoylation of PM proteins. On the other hand, the extraction of FAs from the PM by albumin might decrease the availability of FAs on the cytoplasmic side for activation by acyl-CoA synthetase(s) associated with the PM and their subsequent use by DHHCs to palmitoylate PM proteins. One candidate, ACSL1, appears to directly associate with the FA transporter, FATP1, at the plasma membrane (87). Clear experimental support for one or the other of these hypotheses will have profound implications. In the former case, physical forces within the PM, related to local membrane tension and protein conformation would be key to the regulation palmitoylations at the PM. In the latter case, local turnover and metabolism of FAs will be key to the local control of plasma membrane palmitoylation and the operation of OMDE.

## Acknowledgements

STED microscopy was generously supported by the Peter O’Donnell Jr. Brain Institute NeuroMicroscopy Core. We thank Kate Luby-Phelps of the Quantitative Light Microscopy Core for assistance with STED imaging. We thank Fang-Min Lu for expert technical assistance. Supported by NIH HL119843 and Endowed Professors’ Collaborative Research Support from the Charles Y.C. Pak Foundation. We thank Marcel Mettlen (UTSouthwestern Quantitative Light Microscopy Core Facility) for imaging advice.

## Abbreviations

BBMs: bone marrow macrophages
BP: bromopalmitate
CTB: cholera toxin B
CoA: coenzyme A
CsA: cyclosporine A
FA: fatty acid
β-HB: β-hydroxybutyrate
OMDE: ordered membrane domain endocytosis
PTPs: permeability transition pores
PM: plasma membrane
ROS: reactive oxygen species
TF: transferrin

## Notes

### Competing Interest Statement

The authors have declared no competing interest.

### Summary of Updates

In new experiments we have examined palmitoylation changes in a condition that activates ordered domain endocytosis markedly in cell cultures. In the presence of pyruvate and bicarbonate, we find that ordered domain endocytosis is markedly stimulated and show now that palmitoylations of plasma membrane proteins are markedly increased. These experiments lend new support to the central hypothesis of this article.

