## Supplemental Data for "Constitutive Plasma Membrane Turnover in T-REx293 cells via Ordered Membrane Domain Endocytosis under Mitochondrial Control"

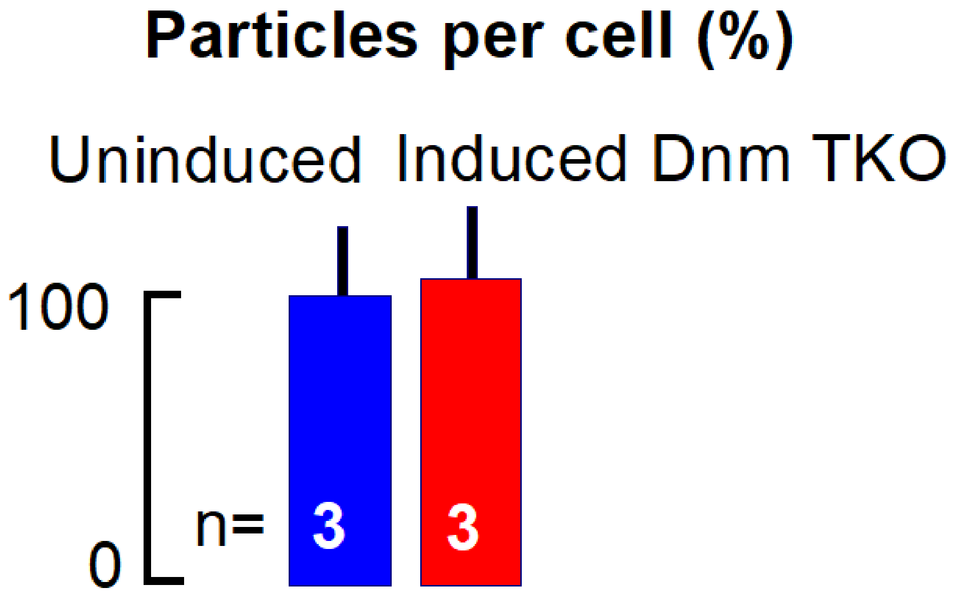


**Supplemental Figure 1.** Mouse embryonic fibroblasts, engineered to allow tamoxifen-induced knockdown of all three dynamin isoforms (1), were cultured and induced as described previously (2). We verify that, as reported, induction of dynamin knockdown did not significantly affect the dextran uptake that occurs over 20 min in serum-, substrate-, and bicarbonate-free conditions.

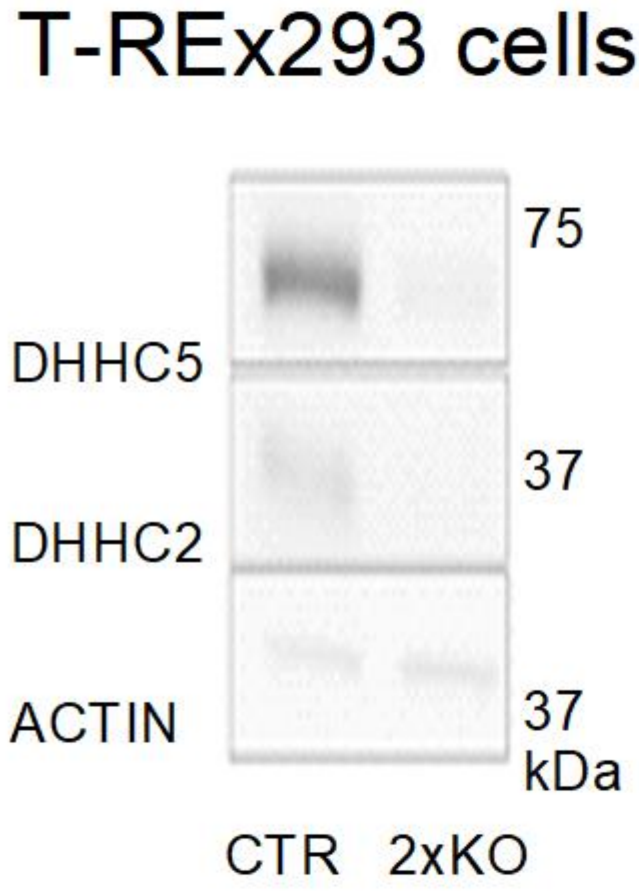


**Supplemental Figure 2.** Western blots of T-REx293 extracts for DHHC5 and DHHC2 from WT cells and cells in which both DHHC isoforms were deleted by CRSPR. Procedures are described in Methods.

Coenzyme A can be rate-limiting for long-chain acyl CoA synthesis.

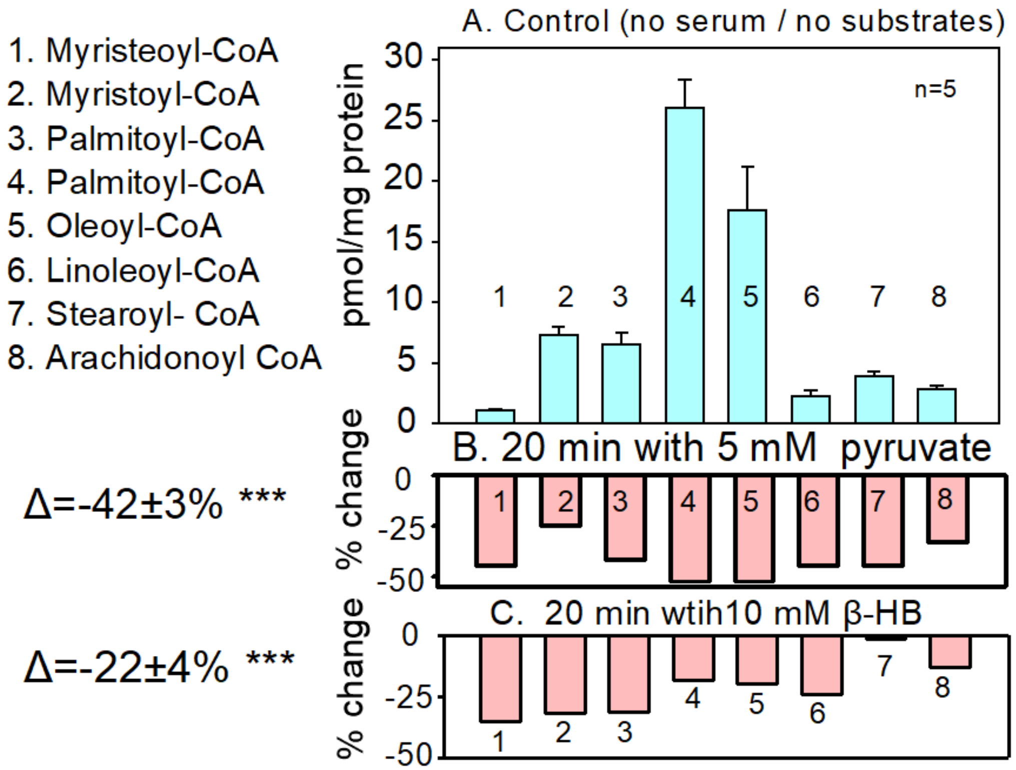


**Supplemental Figure S3.** Analysis of the 8 most prevalent long-chain acyl-CoAs in HEK293 cells. **A.** Concentrations of the indicated acyl CoAs, detemined for 5 dishes of control cells. **B-E.** Relative changes of acyl-CoAs induced by 10 mM of pyruvate, β-HB or carnitine in HEK293 cells after 20 min incubation. Each bar corresponds to the average of 2 to 3 measurements. Average changes of the long-chain acyl-CoA's are given at left. **Methods.** Cells were grown under standard conditions to near confluence. Medium was exchanged for the saline solution employed in serum-, substrate-, and bicarbonate-free conditions for 1 h followed by incubation with the given substrates (10 mM) for 20 min. Acyl-CoAs were quantified by LC-MRM/MS at Creative-Proteomics (Shirley, NY). Briefly, serially diluted calibration solutions containing standard acyl CoAs were prepared in an internal standard (IS) solution of 13C3-malonyl CoA in 80% isopropanol. Each cell pellet sample was thawed on ice. 200 μL of methanol was added. The samples were homogenized on a MM 400 mixer mill with the aid of two metal beads at 30 Hz for 3 min, followed by centrifugation at 21,000 g for 10 min. The clear supernatants were taken out and dried under a nitrogen gas flow. The protein pellets were used for protein assay using a standardized BCA procedure. The dried residues of metabolite extracts were reconstituted in 40 μL of IS solution. 15 μL aliquots of the resultant sample solutions and the calibration solutions were injected to run LC-MRM/MS on a Waters Acquity UPLC system coupled to a Sciex QTRAP 6500 Plus mass spectrometer with negative-ion detection. A 5-cm long C18 UPLC column was used for chromatographic separation, with the use of an ammonium acetate solution and acetonitrile as the mobile phase for binary-solvent gradient elution under an optimized condition. Concentrations of the detected CoAs were calculated with internal standard calibration by interpolating the constructed linear-regression curve with the peak area ratios.

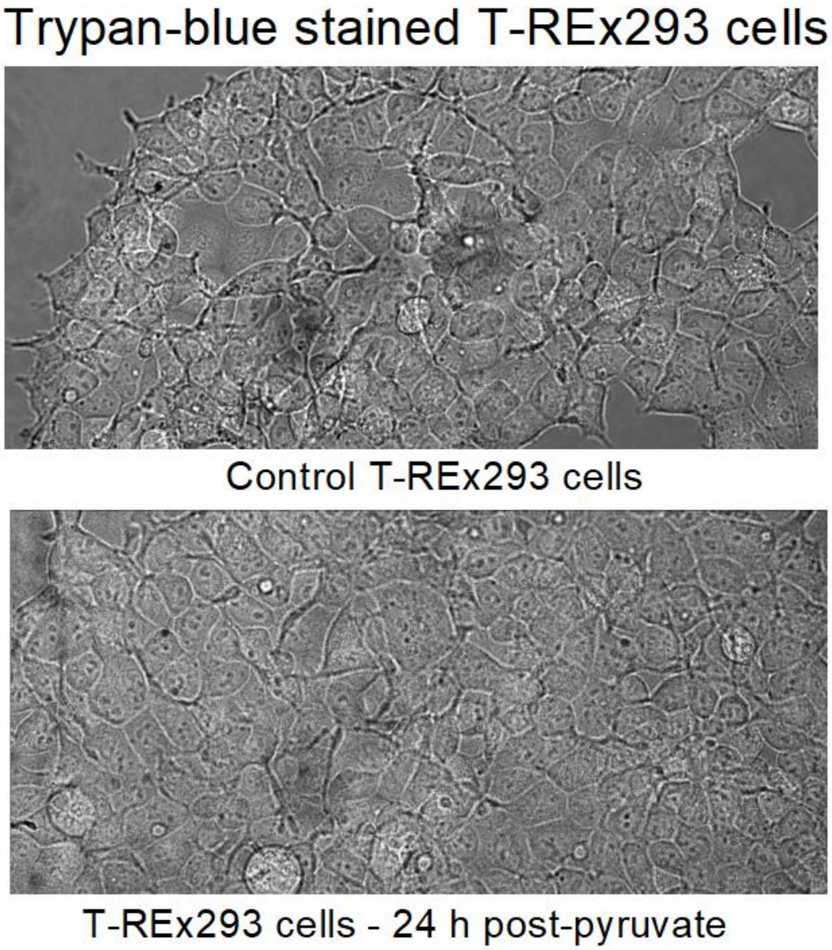


**Supplemental Figure S4.** T-REx293 cell cultures were incubated for 1 h with substrate- and serum-free saline as described in Methods and with Figure 7. Solution was then replaced by the same saline solution containing pyruvate (10 mM) for 15 min and thereafter solution was replaced by standard high glucose DMEM with 10% BFS. Cells were returned to the incubator and 24 h later were subjected to a standard trypan blue protocol to identify dead and dying cells (3). No evidence was found for a loss of cell viability. Thus, the PTPs that may be induced during pyruvate treatment in these experiments do not induce subsequent apoptosis or necrosis.

### Supplemental Figures

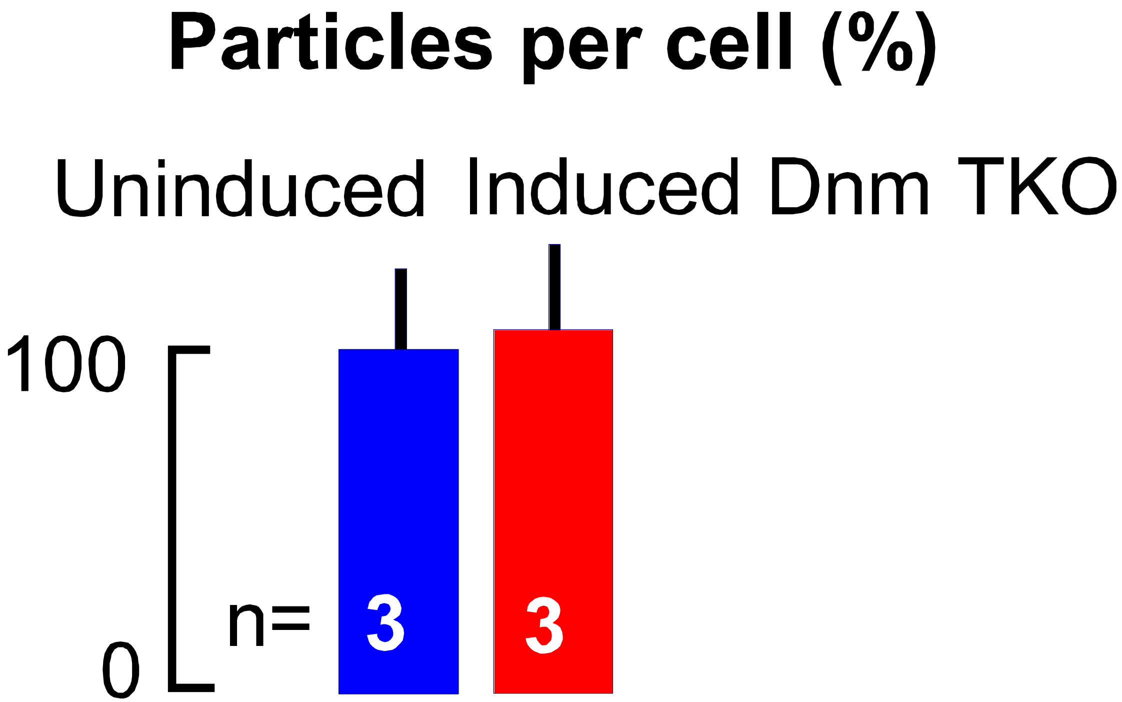


Supplemental Fig. S1

### T-REx293 cells

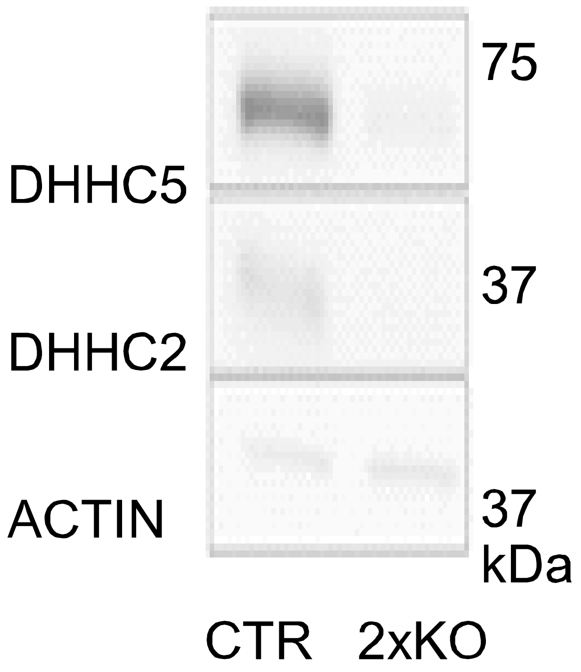


Supplemental Fig. S2

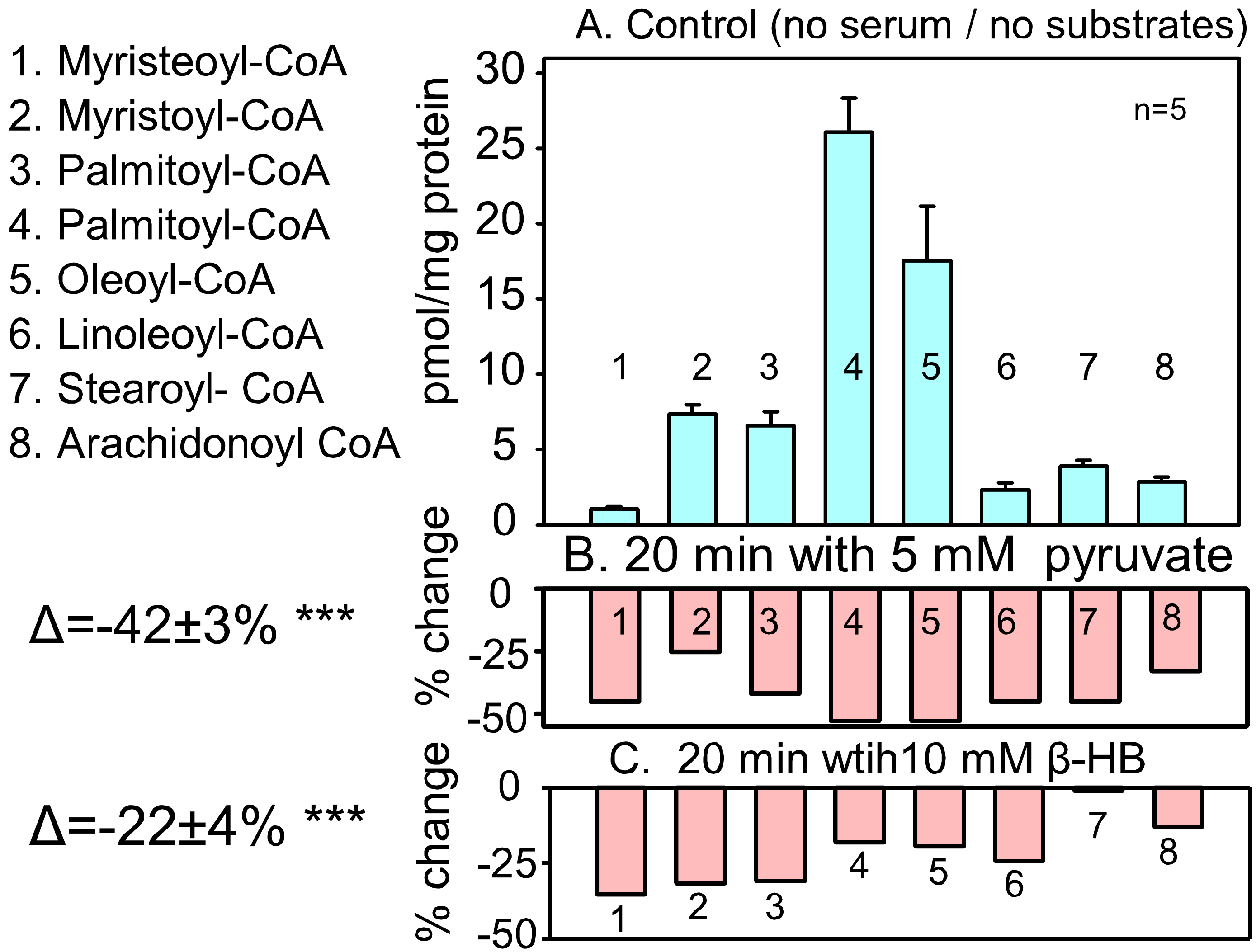


Supplemental Fig. S3

### Trypan-blue stained T-REx293 cells

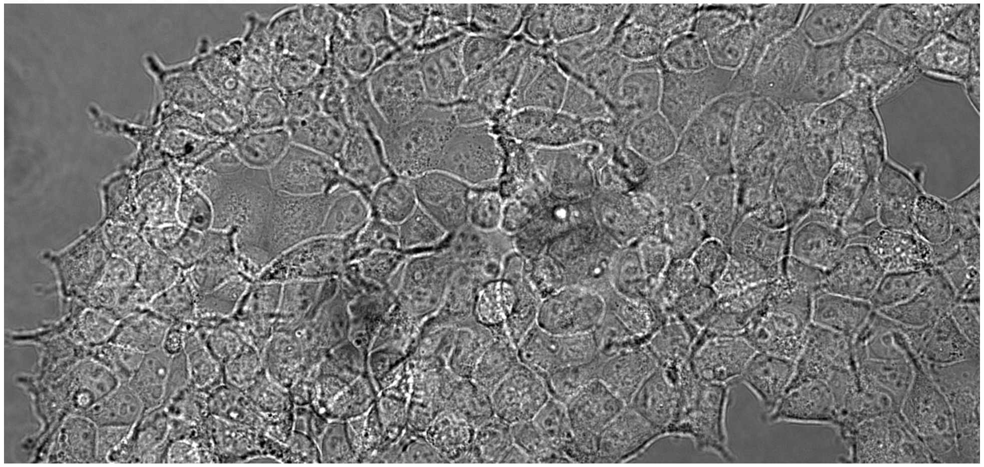

Control T-REx293 cells

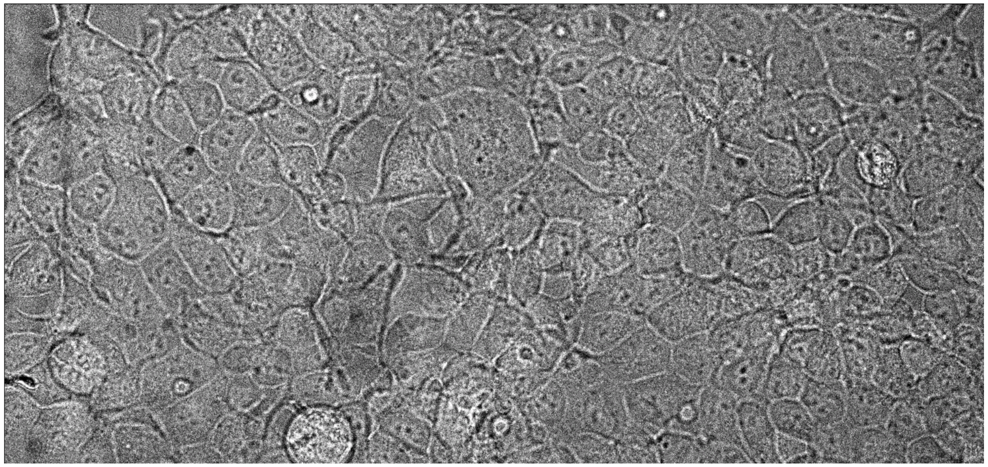

T-REx293 cells - 24 h post-pyruvate

Supplemental Fig. S4
